# Metabolic mapping of the human solute carrier superfamily

**DOI:** 10.1101/2024.09.23.614124

**Authors:** Tabea Wiedmer, Shao Thing Teoh, Eirini Christodoulaki, Gernot Wolf, Chengzhe Tian, Vitaly Sedlyarov, Abigail Jarret, Philipp Leippe, Fabian Frommelt, Alvaro Ingles-Prieto, Sabrina Lindinger, Barbara M. G. Barbosa, Svenja Onstein, Christoph Klimek, Julio Garcia, Iciar Serrano, Daniela Reil, Diana Santacruz, Mary Piotrowski, Stephen Noell, Christoph Bueschl, Huanyu Li, Gamma Chi, Stefan Mereiter, Tiago Oliveira, Josef M. Penninger, David B. Sauer, Claire M. Steppan, Coralie Viollet, Kristaps Klavins, J. Thomas Hannich, Ulrich Goldmann, Giulio Superti-Furga

**Affiliations:** CeMM Research Center for Molecular Medicine of the Austrian Academy of Sciences, 1090 Vienna, Austria; Boehringer Ingelheim Pharma GmbH & Co. KG, 88400 Biberach, Germany; Pfizer Worldwide Research, Development & Medical, Groton CT 06340, USA; Centre for Medicines Discovery, Nuffield Department of Medicine, University of Oxford, Oxford, UK; Department of Laboratory Medicine, Medical University of Vienna, 1090 Vienna, Austria; Institute of Molecular Biotechnology of the Austrian Academy of Sciences (IMBA), 1030 Vienna, Austria; Helmholtz Centre for Infection Research, 38124 Braunschweig, Germany; Department of Medical Genetics, Life Sciences Institute, University of British Columbia, V6T 1Z3 Vancouver, Canada; Center for Physiology and Pharmacology, Medical University of Vienna, 1090 Vienna, Austria

**Keywords:** membrane transporters, metabolism, metabolomics, solute carriers, transcriptomics

## Abstract

Solute carrier (SLC) transporters govern most of the chemical exchange across cellular membranes and are integral to metabolic regulation, which in turn is linked to cellular function and identity. Despite their key role, individual functions of the members of the SLC superfamily were not evaluated systematically. We determined the metabolic and transcriptional profiles upon SLC overexpression in knock-out or wild-type isogenic cell backgrounds. Targeted metabolomics provided a fingerprint of 189 intracellular metabolites, while transcriptomics offered insights into cellular programs modulated by SLC expression. Beyond the metabolic profiles of 102 SLCs directly related to their known substrates, we also identified putative substrates or metabolic pathway connections for 71 SLCs without previously annotated *bona fide* substrates, including SLC45A4 as a new polyamine transporter. By comparing the molecular profiles, we identified functionally related SLC groups, including some with distinct impacts on osmolyte balancing and glycosylation. The assessment of functionally related human genes presented here may serve as a blueprint for other systematic studies of human gene function and supports future investigations into the functional roles of SLCs.

## Introduction

Despite our rapidly increasing familiarity with the human genome and its variant forms, most large-scale studies assessing the genome’s functional properties focus on the contributions of a few genes to specific functional readouts (Kampmann, 2020; Bock *et al*, 2022). There is a need to complement this approach with strategies allowing the comparison of many gene functions, measured with the same readout, in parallel, under controlled conditions. This strategy is particularly suitable for comparing gene families or genes encoding proteins with shared properties since it also generates data for genes that do not emerge in stochastic screens due to redundancy, mildness of phenotype, or specificity of the readout. A particularly worthy area of investigation concerns membrane transporters, responsible for the chemical exchange of biological systems with their environment and the repartition of solutes within biological systems. Not only cellular metabolism depends on the influx and efflux of chemical molecules, but the plethora of physiological processes is chemically integrated and thus also heavily dependent on transporters (Wu *et al*, 2011; Sahoo *et al*, 2014; Alam *et al*, 2023). As the characteristics of the transported molecule(s) are often unknown, and known substrates are chemically very heterogenous, broadly applicable assays have been hard to establish, resulting in many uncharacterized membrane transporters throughout the human genome (Dvorak *et al*, 2021; Saier *et al*, 2016; Fagerberg *et al*, 2010). Therefore, we consider it urgent to better understand this class of proteins (Cesar-Razquin *et al*, 2015; Superti-Furga *et al*, 2020).

Of more than 1,500 genes potentially encoding for transmembrane transporters, we opted to focus on the Solute Carrier (SLC) superfamily as it represents the largest, with ∼450 members (Ye *et al*, 2014; Ferrada & Superti-Furga, 2022; Gyimesi & Hediger, 2022) and is exceptionally asymmetrical in the distribution of knowledge among its members, with a few very well studied and many still uncharacterized (César-Razquin *et al*, 2015; Hediger *et al*, 2013; Oprea *et al*, 2018). The SLC superfamily is divided into >65 families based on their sequence and functional similarity and its members transport a large variety of molecules such as amino acids, inorganic ions, carbohydrates, lipids, neurotransmitters, nucleosides/nucleotides, vitamins, peptides and xenobiotics (Fredriksson *et al*, 2008; Meixner *et al*, 2020, Goldmann et al, accompanying manuscript). SLCs are localized to the plasma membrane or to different intracellular membranes (Meixner *et al*, 2020; Giacomini *et al*, 2022). At the plasma membrane, they control metabolite transport at the cellular level, mediating not only nutrient uptake and waste excretion but also communication and metabolic interaction of the cell with its environment (Pizzagalli *et al*, 2021; Alam *et al*, 2023). Intracellularly localized SLCs are crucial for the creation of different chemical and metabolic environments in the context of compartmentalization of biochemical processes within membrane-surrounded organelles (Bar-Peled & Kory, 2022).

SLCs function as facilitative or secondary active transporters, either equilibrating solute gradients or concentrating solutes by coupling the transport of several substrates through symport or antiport (Colas *et al*, 2016; Drew & Boudker, 2024). The ability to facilitate transport of different molecules or ions along or against a concentration gradient allows transporters to simultaneously control multiple biochemical pathways and the intracellular environment. While some SLCs are very specific for a given substrate, others are promiscuous and able to transport a range of compounds with different chemical structures. This is mirrored from the substrate-centric perspective; while certain compounds are transported by a single SLC, others are substrates for multiple SLCs. Though certain SLCs transport the same substrate, they can still differ in terms of their substrate affinity, specificity, and transport capacity (Meixner *et al*, 2020; Gauthier-Coles *et al*, 2021; Bröer, 2023, accompanying manuscript by Goldmann et al).

Beyond controlling intracellular metabolite concentrations, SLCs influence metabolism by regulating metabolic signals on different levels. This can be at the post-translational level as, for example, glutamine transporters at the plasma membrane regulate mTOR and autophagy, while lysosomal SLC38A9 is involved in the recruitment and regulation of mTORC1, a master regulator of cell growth, by sensing cellular nutrient availability (Nicklin *et al*, 2009; Rebsamen *et al*, 2015; Wang *et al*, 2015). At the transcriptional level, depletion of individual amino acids causes differential transcriptional responses of transporter expression, enabling cells to overcome conditions of nutrient limitation (Rebsamen *et al*, 2022; Chidley *et al*, 2024). For example, nucleoside transporters from SLC families 28, 29 and 35 modulate BRD4-dependent epigenetic states and transcriptional regulation (Li *et al*, 2021). Furthermore, metabolites themselves have regulatory functions and act as important signalling molecules driving feedforward and feedback mechanisms (Cable *et al*, 2021; Baker & Rutter, 2023). SLC-mediated metabolite exchange is not confined to individual cells but plays a critical role in the systemic interactions among distant metabolic organs via the circulatory system. For example, transporters of the SLC2 and SLC5 families are crucial for the uptake and distribution of glucose across the organism (Klip *et al*, 2024).

Understanding how SLCs participate in the regulation of metabolic processes is of therapeutic importance as alteration of transporter function is at the basis of many metabolic perturbations observed both in rare monogenic disorders and common diseases (Zhang *et al*, 2019; Alam *et al*, 2023). Targeting SLCs has already been successful, for example in the treatment of type-2 diabetes by inhibiting SLC5A2 (SGLT2), a sodium-dependent glucose transporter responsible for glucose reabsorption in the kidney (Lin *et al*, 2015). With metabolic reprogramming being a hallmark of cancer (Hanahan, 2022), SLCs are also involved in tumorigenesis, cancer cell growth and metastasis. Cancer cells frequently upregulate glucose and glutamine transporters to meet their increased energetic and biosynthetic requirements (Finley, 2023). SLC-mediated transport also enables adaptation of cancer cells to changes in nutrient availability during the metastatic process (Christen *et al*, 2016; Elia *et al*, 2019; Rinaldi *et al*, 2021; Bian *et al*, 2020). This illustrates how altered metabolism in disease mechanistically converges with metabolite uptake and provides evidence for the therapeutic potential to target SLCs in cancer, though metabolic adaptation is considered a major challenge. Therefore, it is important to understand the interplay between SLC expression and metabolism better.

Addressing the functional complexities of the entire SLC superfamily, in the context of their critical roles in metabolism and the challenges posed by their diverse expression patterns and substrate specificities, necessitates approaches beyond the few traditional cognate assays. Advancements in gene-editing technologies, particularly CRISPR/Cas9, have provided a powerful toolset for selectively manipulating individual SLC genes within a controlled experimental environment. The RESOLUTE consortium (https://re-solute.eu/) was formed to obtain functional data on all SLC transporters encoded in the human genome and shed light on the interface between chemistry and biology that gatekeeps the traffic of chemical substances (Superti-Furga *et al*, 2020). We opted for a comparative, systems-level strategy that incorporates both loss-of-function and gain-of-function methodologies across the entire SLC superfamily and established a collection of genetically modified human cell lines. This methodical approach facilitated a precise assessment of individual SLC roles in steady-state cellular metabolism under standard cell culture conditions. Coordinated in the same large, concerted, effort, involving dozens of academic and industry laboratories, are accompanying studies investigating the global SLC-interactome, systematic genetic interaction mapping and integration of data and knowledge domain in a unified, multimodal landscape (Frommelt and Ladurner et al; Wolf and Leippe et al; Goldmann et al., accompanying manuscripts).

In this study, the basis for the other three studies, we used transcriptomic and metabolomic profiles to establish comparable functional annotations and identify kinship among transporters. This approach leverages two omics techniques to capture a two-dimensional survey of the repercussions of modulating SLC activity. In accordance with the open-access ethos of RESOLUTE, we not only published all generated data sets and developed protocols at open-access databases, but also made most of our created plasmid vectors and cell lines publicly available through non-profit repositories (https://re-solute.eu/resources/reagents). This collection of data sets and reagents (>900 plasmids, >900 sgRNAs), cell lines (>1000) will be an important tool for further elucidating SLC function by the scientific community (**Fig. 1A**).

**Figure 1.**
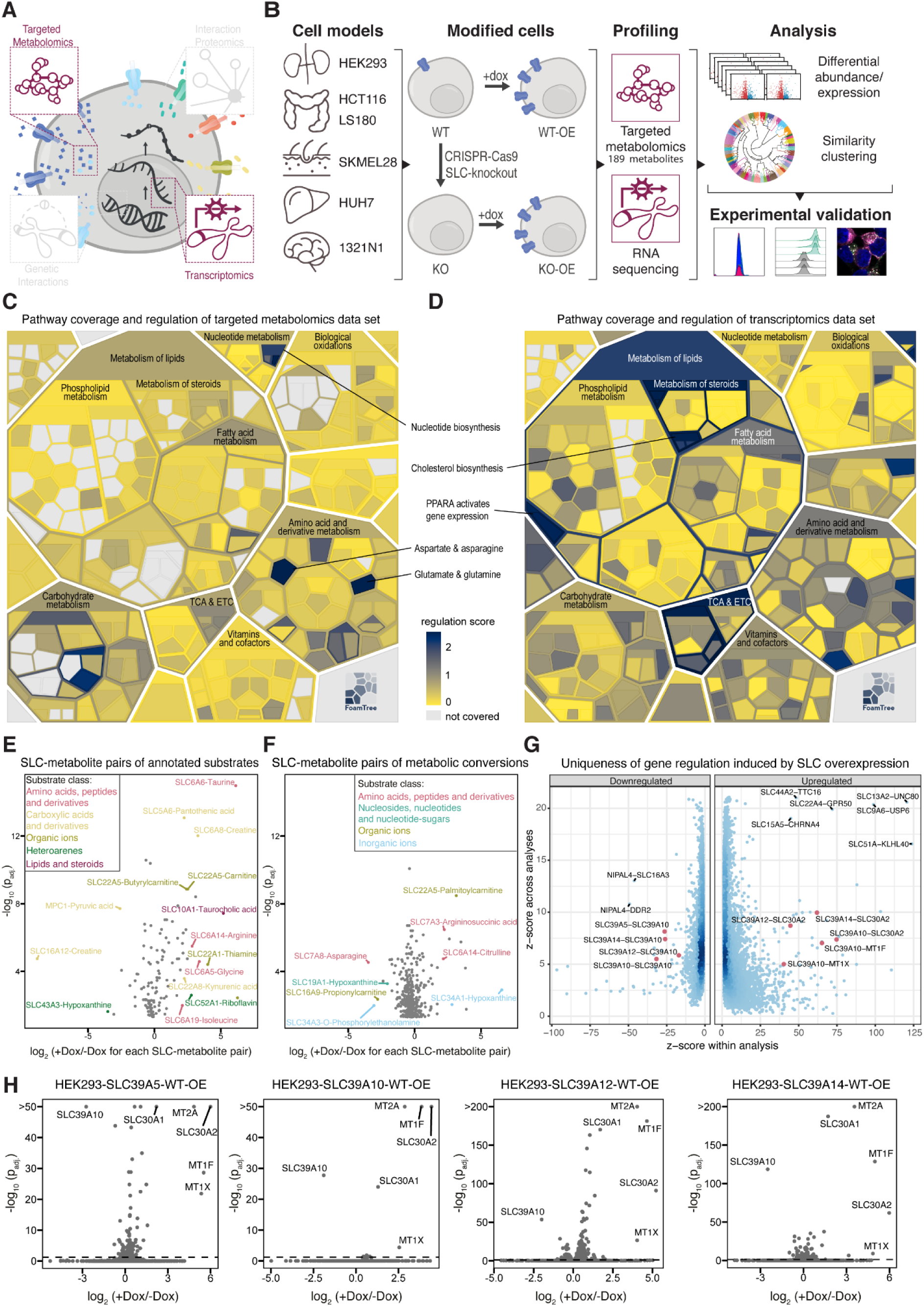
Multi-omic analysis of SLC superfamily-mediated metabolic regulation. **(A)** Overview representation of SLC superfamily-wide data set covered within this RESOLUTE paper collection. **(B)** Schematic representation of workflow for RESOLUTE cell line generation and acquisition and analysis of targeted metabolomics and RNA-sequencing data sets. **(C)** Deregulation frequency of Reactome metabolic pathways based on changed metabolites. For each pathway the number of metabolomics analyses featuring differentially regulated metabolites that were matched to any reaction within the pathway were counted. Significance of the regulation frequency was tested by permutation, shuffling the identities of the quantified metabolites 200 000 times. Significance levels are visualized on a Voronoi treemap of the hierarchical structure of all sub-pathways of human Metabolism pathway in Reactome, to show areas specifically affected by SLC overexpression. (**D)** Deregulation frequency of Reactome metabolic pathways based on deregulated genes. For each pathway the number of transcriptomics analyses featuring differentially regulated genes that were matched to any reaction within the pathway were counted. Significance of the regulation frequency was tested by permutation, shuffling the identities of the quantified genes 200 000 times. Significance levels are visualized on a Voronoi treemap of the hierarchical structure of all sub-pathways of human Metabolism pathway in Reactome, to show areas specifically affected by SLC overexpression. **(E)** Significant SLC- metabolite pairs of SLCs and their annotated substrate. Log2 fold changes and adjusted p-values between +/-doxycycline (dox) of 117 SLC-metabolite pairs of 56 SLCs are visualized. (**F)** Significant SLC-metabolite pairs of SLCs and metabolic conversions of their annotated substrate. Log2 fold changes and adjusted p-values between +dox/-dox of 356 SLC-metabolite pairs of 59 SLCs are visualized. Pairs are colored according to the substrate class of the respective substrate. **(G)** Uniqueness of gene regulation induced by SLC overexpression. Strongly deregulated genes in specific HEK293 WT-OE cell lines (+dox/- dox) were identified by calculation of z-scores of the shrunken log fold change of each gene within a certain analysis and across all analyses. Genes were then further filtered for significance of differential expression (p-adj. < 0.05) and for minimal signal (the gene had to have in one condition at least 50 read counts in both replica). Dots of selected pairs to illustrate the regulation between zinc transporters and metallothioneins are colored in red. (**H)** Differential gene expression analysis between +dox/-dox samples of HEK293-SLC39A5-WT-OE, HEK293-SLC39A10-WT-OE, HEK293-SLC39A12-WT-OE, HEK293-SLC39A14-WT-OE cell lines. P-values were adjusted using the Benjamini-Hochberg correction. The dashed line indicates p- adj. < 0.05.

## Results

### Pipeline for functional profiling of SLC family

For the RESOLUTE project we selected a panel of five human cancer cell lines of different tissue origin (HCT 116/colon, LS180/colon, 1321N1/brain, SK-MEL-28/skin, and Huh-7/liver), cumulatively expressing more than 75% of the SLC superfamily (**EV Fig. 1A**, data from Goldmann et al, accompanying manuscript, ENA Project number PRJNA545487). To study the consequences of SLC function on cellular transcripts and metabolites systematically, we used a controlled cellular system (**Fig. 1B**). For this, we first selected one parental cell line for each individual SLC gene based on its expression pattern across the panel and then generated two independent monoclonal knock-out (KO) cell lines. Subsequently we re-expressed the genetically depleted genes with doxycycline-controlled expression vectors of the cognate codon-optimized cDNA to create two KO-overexpression (KO-OE) cell models per target (details in Methods). An additional set of cell lines with doxycycline-controlled overexpression in wild-type Jump-In™ T-REx™ HEK293 was generated for all SLCs (hereafter referred to as WT-OE). SLC expression in KO-OE and WT-OE cell lines was induced by overnight treatment with doxycycline prior to transcriptomic and metabolomic comparison of uninduced and induced samples. With this two-pronged approach, we aimed to obtain a broad and quantitative picture of the similarities among transporters, as two transporters that elicit similar profiles are likely to be more functionally related than two that are creating dissimilar changes. In some cases, it may also be possible to directly infer the transported substrate from the profile changes.

For transcriptomics, RNA-Seq was performed for 441 SLCs using the WT-OE model only (**EV Fig. 1B**; **Table 1**), covering 99% of the SLC superfamily members and at least one member of each of the 70 SLC families (**EV Fig. 1C**). For targeted metabolomics, we preferably used the KO-OE model to minimize background transporter activity by the endogenous SLC. For SLCs without a KO-OE model available at the time of profiling, we used the HEK293 WT-OE model (**EV Fig. 1B**; **Table 1**). Altogether, targeted metabolomics was performed for 378 SLCs, i.e. 85% of the entire superfamily, covering 67 of the 70 SLC families (**EV Fig. 1D**). We used an ion-pairing reversed-phase liquid chromatography-mass spectrometry method based on a commercially available kit (The Agilent Metabolomics Dynamic MRM Database and Method) as it met our requirements of broad metabolite coverage, robustness of data acquisition, and industry standards for comparability. The method provided reproducible and quantitative measurement of 189 metabolites spanning different compound classes (e.g. nucleotides, amino acids, sugars) and covering most of the central metabolic pathways including glycolysis, TCA cycle, oxidative phosphorylation, pentose phosphate pathway, purine and pyrimidine metabolism, amino acid metabolism, and one-carbon metabolism (**Table 2**).

Differential expression and metabolite abundance between doxycycline-induced and uninduced conditions were analysed for each SLC. Between all SLCs we investigated similarities of metabolomic and transcriptomic profiles, respectively, using hierarchical clustering. We experimentally validated findings from the different computational analyses to showcase approaches to explore the rich data sets, i.e. following up hypotheses either based on profiles of individual SLCs or similarities between groups of SLCs (**Fig. 1B**).

From targeted metabolomics, we observed significant differences between doxycycline-induced and uninduced samples for 282 SLCs (75% of profiled cell lines, metabolites with adjusted p-value < 0.05; **Table 3**). Using a literature-based and manually curated annotation and classification of SLC substrates (Goldmann et al, accompanying manuscript), we grouped SLCs by substrate class and observed that at least 59% of the SLCs in each class showed significant effects, indicating that metabolic changes captured by our method were not biased towards certain SLC substrate classes (**EV Fig. 1E**). From a metabolite-centric perspective, we observed a wide spectrum in the degree and frequency of significant changes across the 189 metabolites included in our method (**EV Fig. 1F, 2A**). We found that 15 metabolites were altered in more than 20% of the cell lines tested. Aconitate, aspartate, and citrate displayed the highest frequency, suggesting that changes in the TCA cycle may be a common effect of perturbing transport-related metabolism. Conversely, 15 metabolites were changed in only one, two or three cell lines and may indicate unique substrates, their metabolic conversions, or uniquely dysregulated metabolic pathways (**EV Fig. 1F**).

Similarly, we assessed which metabolic pathways were altered based on differential metabolite abundance and gene expression upon SLC overexpression. Out of a total of 328 metabolic pathways in the Reactome database (https://reactome.org/PathwayBrowser/#/R-HSA-1430728; (Milacic *et al*, 2024)), 257 and 308 pathways were differentially regulated based on metabolite abundance and gene expression, respectively, across all profiled SLCs (**Fig. 1C, 1D, Table 4**). Therefore, our approach enabled the interrogation of a considerable portion of the metabolic space with the two data modalities complementing each other. Permutation analysis and subsequent enrichment test showed that in the metabolomics data set, metabolites in carbohydrate, nucleotide, and amino acid metabolism were most frequently deregulated while in the transcriptomics data set genes in lipid metabolism, the TCA cycle, and the electron transport chain, were most frequently deregulated (**Fig. 1C, 1D**). This was in line with the analysis at the metabolite level (**EV Fig. 1F**) and indicated that substrate transport often caused perturbation of energy metabolism.

Next, we searched the differential metabolite abundance profiles for known substrates of SLCs to validate that the workflow could also assess directly transported substrates. 279 of the 378 profiled SLCs had one or more annotated substrates and for 141 of them, our targeted metabolomics method covered at least one substrate. For 57 of those SLCs (40%) from different substrate classes, we identified significant changes of annotated cognate substrates, such as SLC5A6:pantothenic acid, SLC6A8:creatine, SLC22A5:carnitine and SLC6A6:taurine pairs, for which we observed increased substrate levels. Conversely, we found decreased cellular levels of pyruvate upon overexpression of MPC1, consistent with pyruvate utilization in mitochondria **(Fig. 1E)**. Since altered transport not only affects steady-state levels of transported substrates but also their metabolic derivatives, we expanded the analysis to metabolic conversions, i.e. metabolites which are either educts or products of reactions involving annotated SLC substrates. For this, we mapped annotated SLC substrates to Reactome pathway reactions and considered adjacent metabolites covered by our targeted metabolomics method. Of the 279 substrate-annotated SLCs included in this study, our method covered the measurement of such metabolic conversions for 233 SLCs. For 82 of those SLCs (35%), we found the levels of either an educt or a product of a metabolic reaction involving at least one of their annotated substrates significantly changed (**Fig. 1F)**. The frequency of significant changes was significantly higher when the metabolite was either an annotated substrate or a metabolic conversion compared to the regulation frequency of all metabolites across all cell lines (Fisher’s exact test, both p<0.0001) (**EV Fig. 1G**). Although our targeted metabolomics approach mainly focused on molecular profiling of the functional consequences downstream of SLC expression, these results showed that our approach could also measure direct or closely proximal substrate abundance changes for a proportion of SLCs (102 out of 279 profiled SLCs with substrate annotation).

Arguably, the transcriptional response to SLC overexpression is less suitable for the identification of direct chemical substrates of the SLCs under investigation. However, we reasoned that gene expression changes should be mainly elicited by the activity of the transporter under investigation, i.e. by changing the concentration of specific metabolites and ions. Importantly, the transcriptional profile is quantitative and robust, covers several thousand genes involved in virtually all cellular functions, has been the workhorse for differential molecular responses of cells to perturbations for decades, and is one of the most interoperable phenotypic descriptor of cell states (Lamb *et al*, 2006; Niepel *et al*, 2017; Keenan *et al*, 2018; Subramanian *et al*, 2017).

In our transcriptomic data set many deregulated genes were affected in multiple different cell lines, suggesting that they may reflect transport-independent effects, such as a general cellular stress response caused by high transgene expression levels. Nevertheless, the approach also identified genes that are uniquely deregulated through the expression of specific SLCs (**Fig. 1G**; **Table 5**). For example, the genes for the metallothionein family members MT1F, MT1X and MT2A, that are known to protect against metal toxicity by binding heavy metals such as zinc and copper (Chen *et al*, 2024), were upregulated in cell lines overexpressing one of the zinc importers SLC39A5, SLC39A10, SLC39A12 or SLC39A14 (**Fig. 1H**). Additionally, two zinc exporters, SLC30A1 and SLC30A2, were upregulated in these cell lines, likely to counter the excessive zinc influx, while the zinc importer SLC39A10 was downregulated (**Fig. 1H**). Altogether, these data indicated that the increased metal import activity in these cells triggered a transcriptional response aimed at lowering the concentration of unbound intracellular metals. Importantly, this exemplified how transcriptomics may inform on transport of substrates not covered by the targeted metabolomics method. Further, it suggested how superfamily-wide systematic perturbation of SLCs may provide insights into function of individual SLCs through comparable transcriptional responses, conceptually validating the approach.

### Insights to SLC function by metabolite-SLC pairs

To identify strong metabolomic changes that could potentially lead to new substrate identification in validation studies, we used a rationale based on the benchmarking data set of annotated substrates and metabolic conversions for SLCs as described above. We grouped all measured SLC-metabolite relationships, that we call ‘pairs’, into four categories: i) ‘expected’ pairs of SLCs with their annotated substrates (‘annotated substrate’); ii) ‘expected’ pairs of SLCs and metabolic conversions of their substrate, i.e. an educt or product of a reaction with the substrate as described previously (‘metabolic conversion’); iii) ‘novel’ SLC-metabolite pairs, consisting of metabolite hits not matching annotated substrates or their immediate metabolic conversions (‘novel for non-orphan SLC’); iv) metabolite hits for orphan SLCs (‘novel for orphan SLC’) (**Fig. 2A**; **Table 3**). We compared the magnitude of change upon differential abundance analysis across the four categories and found that changes in metabolite levels of SLC-annotated substrate pairs were significantly larger than those of SLC-metabolic conversion pairs (mean log2 fold changes, Kruskal-Wallis test, p<0.0001). This indicated a tendency for direct substrates to show larger abundance changes upon SLC overexpression, a finding that is not surprising as metabolic conversion may be multidirectional and diffused. This also suggested that novel SLC-metabolite pairs with a relatively large log2 fold change may imply novel putative substrates for the respective SLCs (**Fig. 2A**). To that end, the data set was explored on a per-metabolite basis, which illustrated on the one hand all SLC-metabolite pairs divided per metabolite and, on the other hand, the distribution of magnitude of change as well as the SLCs causing the largest changes (**EV Fig. 2A**).

**Figure 2.**
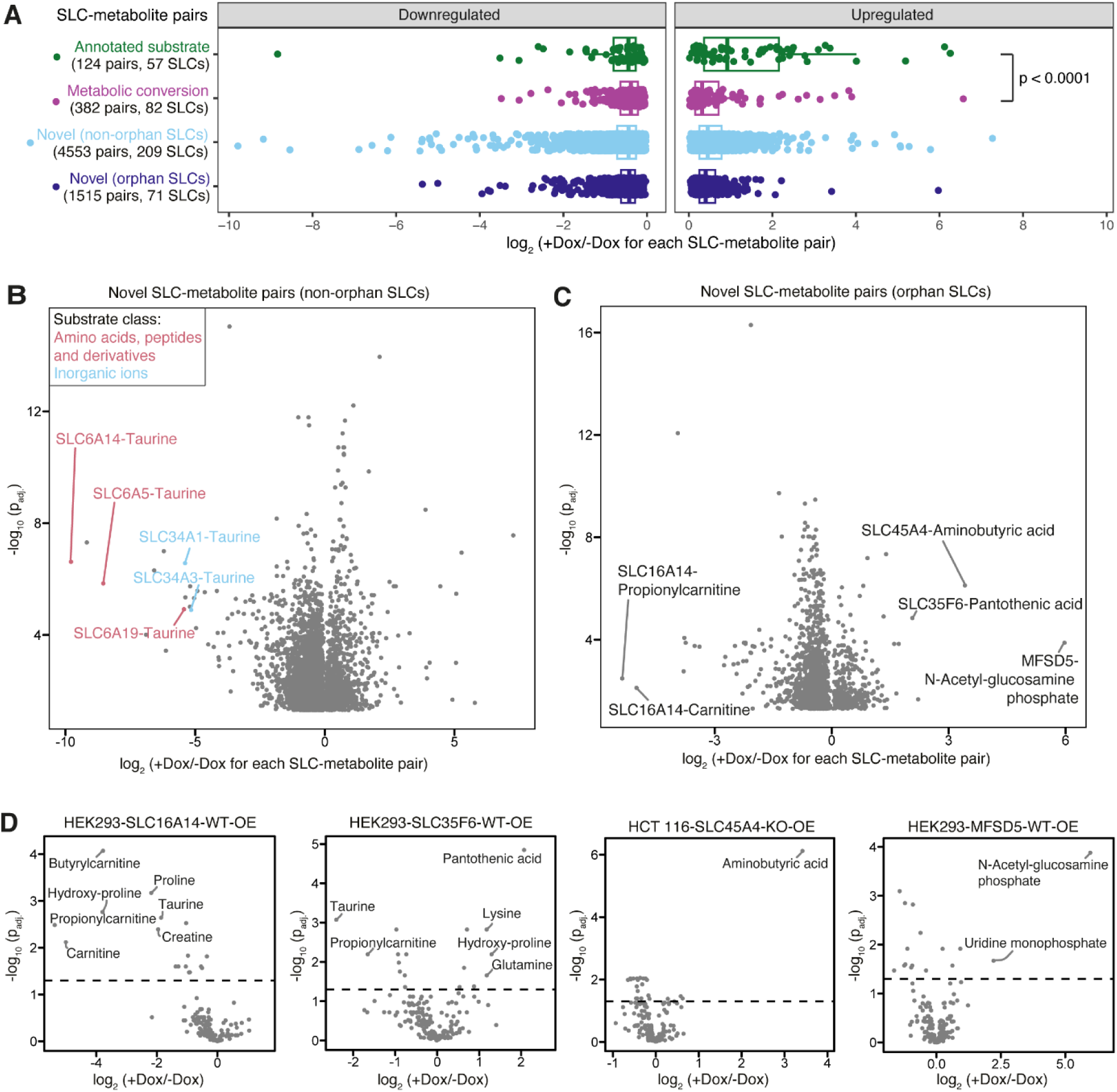
Deregulated SLC-metabolite pairs illuminate known and novel functional relationships. **(A)** Boxplot of all SLC- metabolite pairs divided into categories ‘annotated substrate’, ‘metabolic conversion’, ‘novel (non-orphan SLCs)’, ‘novel (orphan SLCs)’. Number of pairs and respective number of SLCs are indicated per category. Kruskal-Willis test was performed between all categories and the p-value for the significant difference indicated (only between increased metabolite levels of ‘annotated substrate’ and ‘metabolic conversion’ categories (p < 0.0001). **(B)** Significant novel SLC-metabolite pairs for SLCs with an annotated substrate. Respective metabolites are not substrates or metabolic conversions. Pairs are colored according to the substrate class of the respective substrate. **(C)** Significant novel SLC-metabolite pairs for orphan SLCs. **(D)** Differential metabolite abundance analysis between +/-doxycycline (dox) samples of HEK293-SLC16A14-WT-OE, HEK293-SLC35F6-WT-OE, HCT116-SLC45A4-KO-OE, HEK293-MFSD5-KO-OE cell lines. P-values were adjusted using the Benjamini-Hochberg correction. The dashed line indicates p-adj. < 0.05.

For 209 SLCs with annotated substrates in the knowledge domain, we found >4,500 SLC-metabolite pairs significantly changed that had not previously been annotated in connection with these SLCs (category iii, **Fig. 2A**). Taurine was among the most strongly altered metabolites with a more than 5- fold decrease upon expression of five SLCs: SLC6A5, SLC6A14, SLC6A19, SLC34A1 and SLC34A3. While these SLCs had diverse annotated substrates, such as different amino acids and phosphate, all of them couple substrate transport to sodium (Na^+^) (**Fig. 2B**, Goldmann et al, accompanying manuscript). This suggested that SLC-induced taurine reduction may have been related to osmotic disbalance and/or sodium coupling.

Lastly, we found > 1,500 SLC-metabolite pairs for 71 orphan SLCs (category iv, **Fig. 2A**). Among these, the strongest changes were observed for SLC45A4:gamma aminobutyric acid (GABA), SLC35F6:pantothenic acid, MFSD5:n-acetylglucosamine phosphate, SLC16A14:proprionylcarnitine and SLC16A14:carnitine (**Fig. 2C**). Therefore, these SLCs were considered for further experimental validation. As a higher number of changed metabolites may point towards a heavily perturbed cellular homeostasis, we compared the metabolic profiles of SLC45A4, SLC35F6, MFSD5 and SLC16A14, that showed a more than 2-fold change of one, six, eight, and eleven metabolites, respectively (**Fig. 2D**). Evaluating the frequency of significant changes in the respective metabolites across all profiled SLCs, GABA was among the least frequently changed metabolites (changed only in six cell lines), while pantothenic acid, n-acetylglucosamine phosphate, propionylcarnitine and carnitine showed significant changes only in 3% to 25% of cell lines (**EV Fig. 1F**). Based on the strong differential change and the unique metabolite profile, we focused on the SLC45A4:GABA pair for further validation.

### SLC45A4 mediates the export of polyamines

Immunofluorescence analysis using the HA-tag in HCT116 SLC45A4 KO-OE cells, showed that SLC45A4 localized to the plasma membrane (**EV Fig. 3A**). GABA supplementation to culture media caused a time-dependent increase in intracellular GABA concentration of HCT116 cells overexpressing the GABA transporter SLC6A1, but not SLC45A4, ruling out direct uptake as the mechanism of SLC45A4-mediated GABA increase (**EV Fig. 3B)**. Since the heterologous expression of mouse SLC45A4 in yeast cells has been reported to mediate sucrose uptake (Bartölke et al. 2014), we also measured sucrose uptake but did not observe increased uptake in SLC45A4 overexpressing cells (**EV Fig. 3C).** We next examined the metabolic pathways involved in intracellular GABA production, i.e. through the GABA shunt via direct conversion of glutamic acid via glutamate decarboxylase, encoded by *GAD1* and *GAD2*, or through the polyamine synthesis pathway from putrescine by diamine oxidase (DAO), encoded by the *AOC1* gene (Amine Oxidase, Copper-containing, 1) (**Fig. 3A**). Using stable isotope labelling, we probed the contribution of these pathways to GABA production in our cellular system. Culturing HCT116 SLC45A4 KO-OE cells in ^13^C^15^N-glutamic acid did not lead to the production of labelled GABA (**EV Fig. 3D**). However, GABA was labelled upon culturing the cells in media supplemented with ^13^C^15^N-arginine, a precursor for polyamines. Additionally, we observed substantial labelling of ornithine and the polyamines putrescine, spermidine and spermine, supporting the notion that GABA may originate from putrescine (**Fig. 3B, upper row**). Labelled putrescine and spermidine levels decreased intracellularly and were elevated in supernatants of doxycycline-induced cells, suggestive of increased export of polyamines (**Fig. 3B, lower row**). To confirm that increased GABA derived from putrescine, cells were treated with difluoromethylornithine (DFMO), an inhibitor of ornithine decarboxylase (ODC1), the enzyme that produces putrescine from ornithine. ODC1 inhibition caused a stark depletion in intracellular putrescine levels and blocked SLC45A4-mediated GABA accumulation (**EV Fig. 3E**). Hence, SLC45A4 appeared to mediate an increase in putrescine-derived GABA, justifying an in-depth investigation of the pathway from putrescine to GABA.

**Figure 3.**
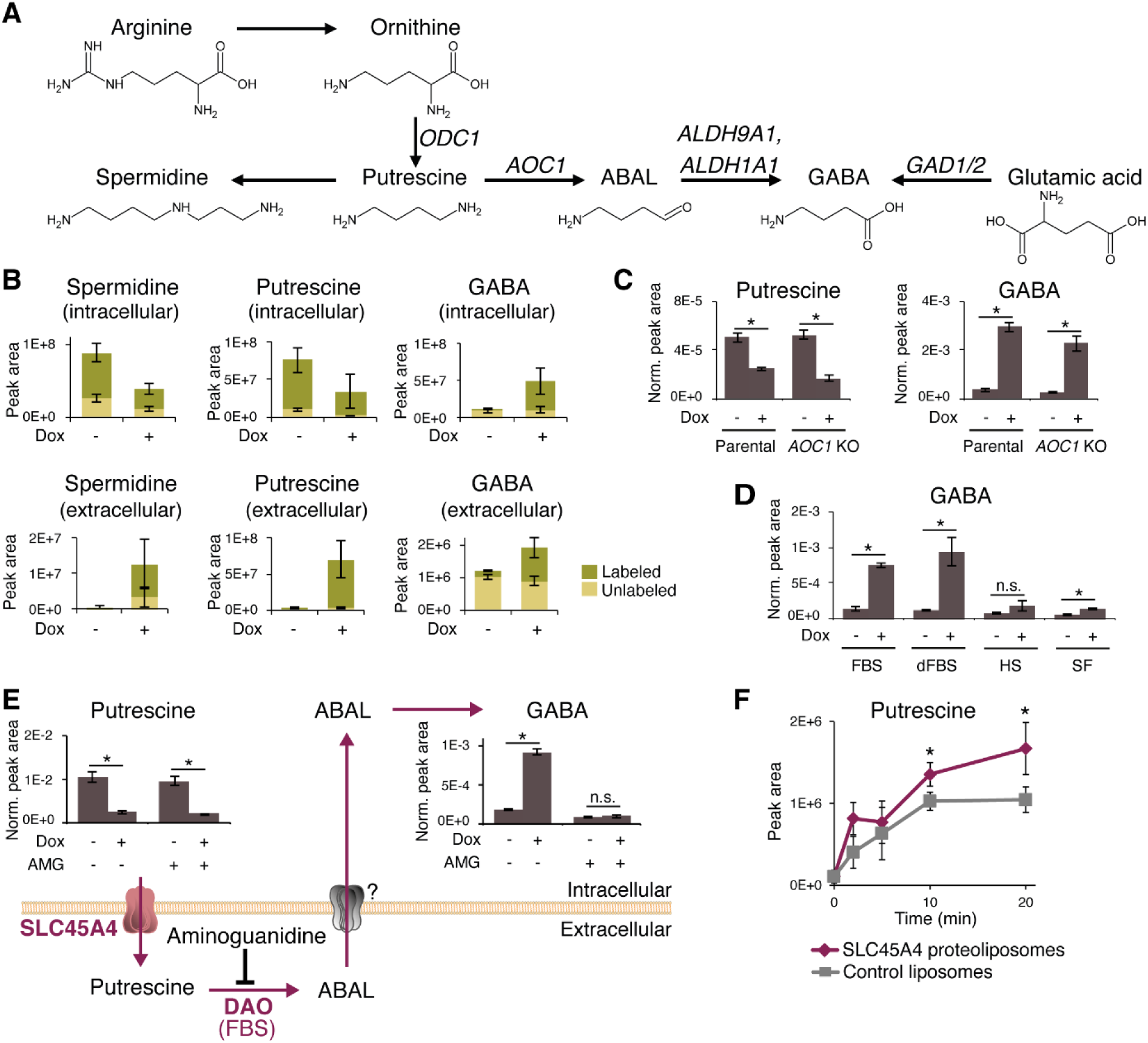
The orphan SLC45A4 is a novel putrescine exporter. **(A)** Schematic representation of metabolic pathways that produce GABA. **(B)** Labeled (from ^13^C^15^N-arginine) and unlabeled abundances of intracellular vs extracellular metabolites for HCT116 SLC45A4 KO-OE cells +/- 24 h doxycycline (Dox) induction. Bar heights represent means, error bars represent s.d. (n=6). **(C)** Intracellular putrescine and GABA levels in HCT116 SLC45A4 KO-OE cells with and without AOC1 KO +/- 24 h Dox induction. Bar heights represent means, error bars represent s.d. (n=3). **(D)** Intracellular GABA levels in HCT116 SLC45A4 KO- OE cells cultured in regular FBS-containing media, media with dialyzed FBS (dFBS), media with horse serum (HS), or serum- free media (SF) +/- 9 h Dox induction. Bar heights represent means, error bars represent s.d. (n=3). **(E)** Intracellular putrescine and GABA levels in HCT116 SLC45A4 KO-OE cells with and without DAO inhibition by aminoguanidine (AMG) +/- 9 h Dox induction. Bar heights represent means, error bars represent s.d. (n=3). Purple arrows show the proposed pathway leading to increased intracellular GABA accumulation in SLC45A4 overexpression cells: SLC45A4 mediates export of putrescine into the extracellular space, where it is then acted upon by serum-derived DAO activity to generate ABAL for GABA production. **(F)** Putrescine quantification in SEC eluates of SLC45A4 proteoliposomes vs control liposomes over different uptake durations. Data points represent means, error bars represent s.d. (n=3). Where presented, asterisks (*) and n.s. on significance bars indicate p < 0.05 and p ≥ 0.05 (Student’s t-test) respectively.

AOC1, the human DAO responsible for converting putrescine to 4-aminobutyraldehyde (ABAL) for subsequent oxidation to GABA, is not expressed in HCT116 cells (data from Goldmann et al, accompanying manuscript; ENA Project number PRJNA545487). In line with this, AOC1 KO did not affect putrescine export or GABA accumulation (**Fig. 3C**). Since fetal bovine serum (FBS), a component of regular cell culture media, contains DAO activity (Gahl & Pitot, 1979), we hypothesized that the putrescine to GABA conversion occurred through an extracellular pathway and that serum-derived DAO activity converted extracellular putrescine to ABAL. We cultured HCT116 SLC45A4 KO-OE cells in different media serum conditions. Substitution of regular FBS with dialyzed FBS (dFBS), from which small molecules have been excluded, had no effect on SLC45A4-mediated GABA production. Conversely, SLC45A4 KO-OE cells cultured in horse serum (HS), which has low DAO activity (Kunimoto *et al*, 1985; Niskanen & Wharton, 1987), or serum-free media (SF) failed to accumulate GABA (**Fig. 3D**) while depletion of intracellular putrescine was unaffected (**EV Fig. 3F**). This suggested that SLC45A4 exported putrescine and spermidine, while serum DAO activity converted exported putrescine into ABAL for subsequent GABA synthesis. Indeed, DAO inhibition by aminoguanidine (Caron *et al*, 1988) blocked GABA accumulation in SLC45A4 KO-OE cells (**Fig. 3E**). Additionally, while cells cultured in HS-supplemented culture media accumulated significantly less GABA than in FBS, supplementation of porcine DAO to the media greatly increased the amount of GABA production for both FBS and HS conditions (**EV Fig. 3G**), indicating that extracellular DAO activity was a limiting factor in the production of GABA from putrescine. To assess whether SLC45A4 directly transports putrescine, we performed an uptake assay using liposomes reconstituted with purified SLC45A4 protein (**EV Fig. 3H**). Putrescine levels measured by LC-MS analysis increased over time upon incubation in assay buffer and were significantly higher in SLC45A4 proteoliposomes relative to control liposomes (**Fig. 3F; EV Fig. 3I**). Hence, our data indicate that SLC45A4 directly transports putrescine.

### Similarity of omics profiles indicates functional relationships between SLCs

While the combined transcriptomics and metabolomics data allow for generation of a large number of hypotheses on individual SLCs worth further investigation, we next explored the potential for more general observations. We expanded the analysis from individual SLC profiles to an SLC superfamily-wide analysis, aiming to identify metabolic and transcriptomic patterns among SLCs that would facilitate broad functional assignment through profile similarity. Employing hierarchical clustering, we grouped all overexpression cell lines based on the differential response of 14,187 genes and 142 metabolites between induced and uninduced samples. We selected the number of clusters based on the mean silhouette width (Rousseeuw, 1987), aiming for an average cluster size of 5-10 cell lines (**EV Figs. 4A, 5A**).

**Figure 4.**
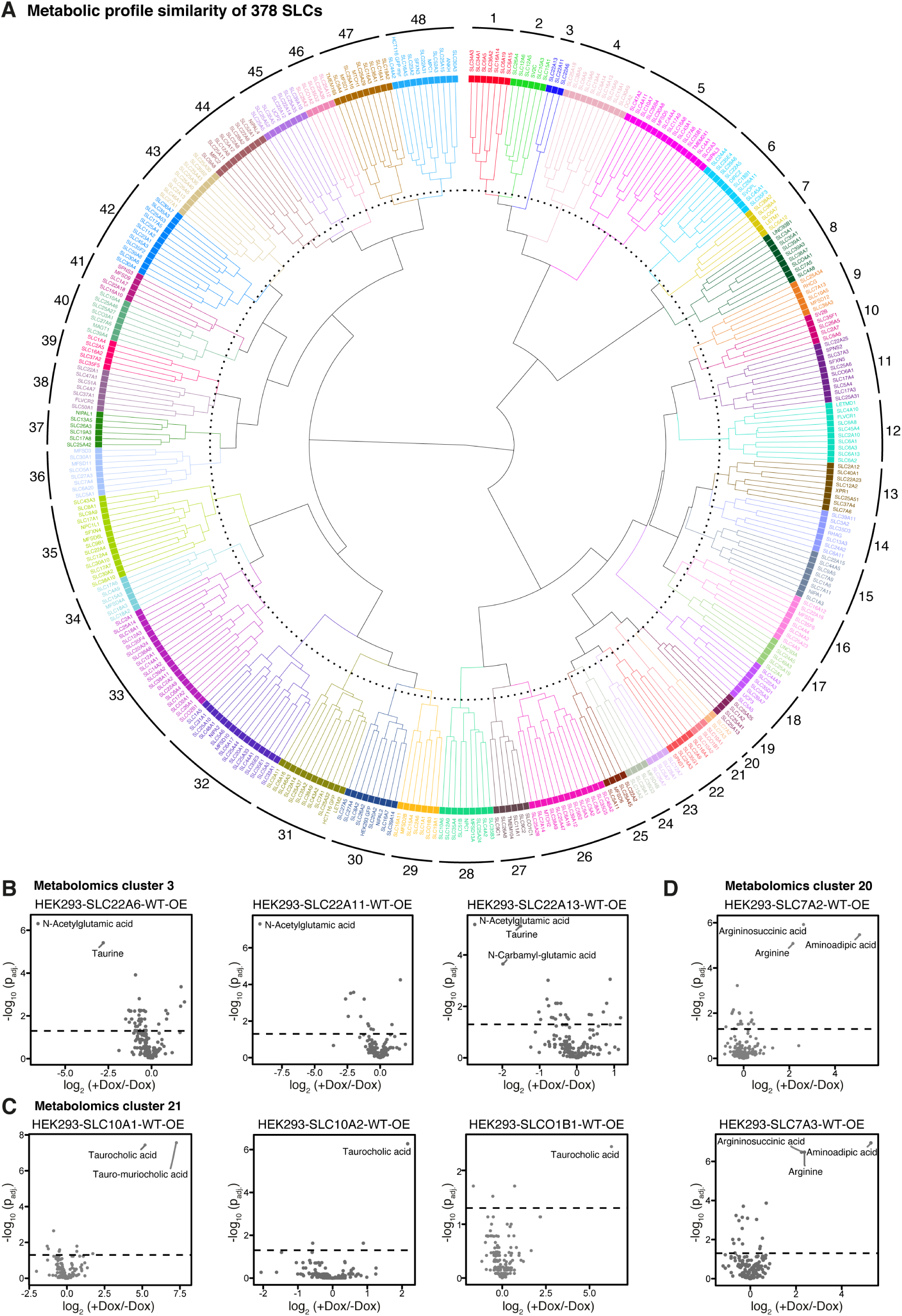
Similarity clustering of metabolomic profiles reveals known and novel SLC functional relationships. **(A)** Dendrogram of hierarchical clustering based on the Euclidean distance matrix of normalized metabolic profiles of 381 cell lines (378 SLC cell lines and 3 GFP cell lines). **(B)** Differential metabolite abundance analysis upon +/- doxycycline (Dox) induction of HEK293-SLC22A6-WT-OE, HEK293-SLC22A11-WT-OE and HEK293-SLC22A13-WT-OE cell lines. **(C)** Differential metabolite abundance analysis upon +/- Dox induction of HEK293-SLC10A1-WT-OE, HEK293-SLC10A2-WT-OE and HEK293- SLCO1B1-WT-OE cell lines. **(D)** Differential metabolite abundance analysis upon +/- Dox induction of HEK293-SLC7A2-WT-OE and HEK293-SLC7A3-WT-OE cell lines. P-values were adjusted using the Benjamini-Hochberg correction. The dashed line indicates p-adj. < 0.05.

**Figure 5.**
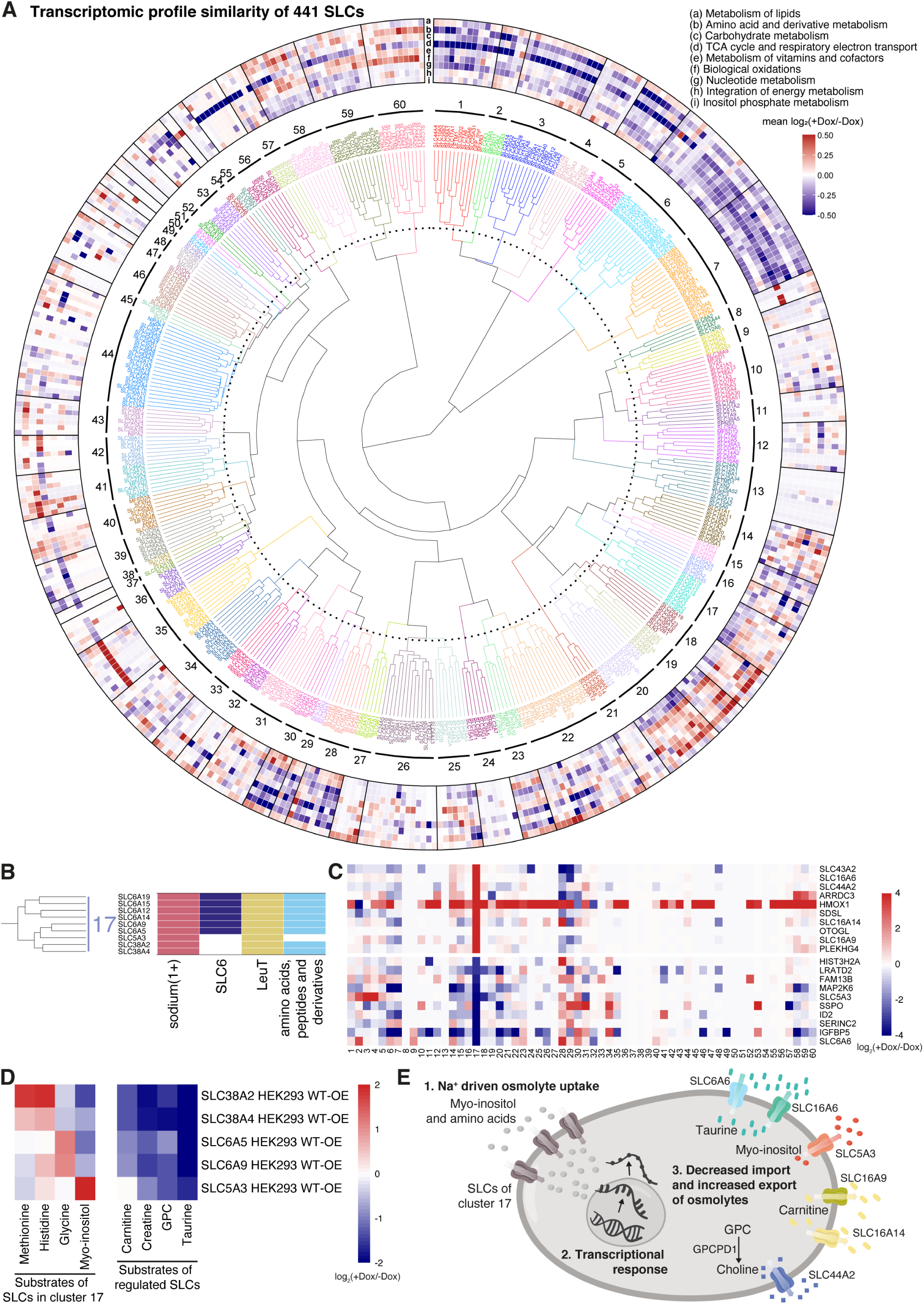
Similarity clustering of transcriptomic profiles identifies differential metabolic impacts of SLC functional groups. **(A)** Dendrogram of hierarchical clustering based on the Euclidean distance matrix of standard normalized transcriptional profiles of 450 analyses (441 SLC and 3 GFP cell lines; for 5 SLCs two analyses with different induction times). The outer ring displays mean log2 fold-change of metabolic genes in each of 9 top-level Reactome pathways. **(B)** Significantly enriched SLC functional properties for cluster 17 (Fisher’s test p < 0.2). **(C)** Heatmap showing doxycycline (Dox) vs uninduced average log2 fold changes in each cluster, for the 20 most deregulated genes in cluster 17 compared to other clusters. Cluster numbers are indicated on x-axis. **(D)** Heatmap of Dox vs uninduced log2 fold changes of known substrates of SLCs in cluster 17 as well as metabolites expected to be perturbed by commonly deregulated SLCs and metabolic enzymes in the cluster. **(E)** Illustration of osmolyte uptake and transcriptional response to balance intracellular osmolyte concentrations and decrease osmotic pressure.

Metabolomic profiles of 378 SLC and three GFP-expressing cell lines resulted in 48 clusters, based on their similarity in differential abundance of the 142 metabolites upon overexpression (**Fig. 4A**; **Table 6**). Using a heatmap of the average metabolic profile per cluster, we identified key factors driving the similarities between clustered SLCs. This analysis led us to further explore selected clusters, their individual members, and the associated metabolite changes (**EV Fig. 2B**). For instance, we found cluster 3 consisting of SLC22A6, SLC22A11, and SLC22A13, which all strongly reduced N-acetylglutamic acid levels (**Fig. 4B**). All three SLC22 members are annotated to transport urate and increased urate uptake is anticipated to inhibit N-acetylglutamic acid production due to urate’s role as a N-acetylglutamate synthase inhibitor (Nissim *et al*, 2011), likely to explain the observed phenotype. Cluster 21 was characterized by the increase of taurocholic acid, an annotated substrate for all its members, SLC10A1, SLC10A2 and SLCO1B1 (**Fig. 4C**). Cluster 20 with SLC7A2 and SLC7A3 as members presented an example in which we observed similarity due to the differential signal of both the annotated substrate arginine as well as the metabolic products argininosuccinic acid and aminoadipic acid, which are downstream of arginine and lysine (another annotated substrate), respectively (**Fig. 4D**). Furthermore, hierarchical clustering of cluster-averaged metabolite profiles revealed co-regulation of several metabolite groups including nucleotide monophosphates, nucleotide triphosphates, and amino acids (**EV Fig. 4B**). For instance, the SLCs within cluster 1 (SLC34A3, SLC34A1, SLC6A5, SLC36A2, SLC16A14, SLC6A19, SLC6A15) showed an upregulation of a group of metabolites consisting of isoleucine, tryptophan, tyrosine, histidine, leucine, citrulline, valine and methionine. Besides the upregulation of several amino acids, cluster 1 was characterized by a strong downregulation of taurine that likely stemmed from osmotic imbalance due to sodium-coupled transport, which had been reported for all SLCs in the cluster except SLC36A2 and SLC16A14 (Goldmann et al, accompanying manuscript). Another commonality between the amino acid transporters of this cluster was their transport of glycine and/or proline, that may present an interesting starting point to study the regulatory relationships between amino acid transporters. Generally, these observations suggested that the grouping of SLCs by the gain-of-function-elicited metabolic fingerprint allowed for the identification of functional relationships.

In analogy to the analysis based on metabolites, we clustered SLCs based on their transcriptional profile. 447 SLC and three GFP cell lines were clustered into 60 groups based on their similarity in the differential expression profile of 14,187 genes (**Fig. 5A**; **Table 7**). To obtain a coarse overview of metabolic effects driven by each transcriptomic cluster, we calculated mean expression changes of genes associated with each Reactome pathway and examined the 9 top-level pathways (**Fig. 5A, outer ring; Table 8**). Clusters 1 to 7 showed broad deregulation of metabolism, with a general trend of downregulation. Other clusters had strong, specific signals for a single pathway, e.g. clusters 8, 35, 56 for biological oxidations and the TCA cycle. On the other hand, certain clusters did not appear to have strongly perturbed gene expression in any of the broad categories of metabolism such as for example clusters 9 and 13.

Next, we evaluated each cluster for enrichment of shared functional properties of the overexpressed SLCs such as SLC family membership, structural fold, substrate class or subcellular localization (**Appendix Figure S1)**. Two clusters stood out with significant overrepresentation of four functional SLC properties each: cluster 17 (SLC6A19, SLC6A15, SLC6A12, SLC6A14, SLC6A9, SLC6A5, SLC5A3, SLC38A2 and SLC38A4) and cluster 6 (17 SLCs from diverse families) (**EV Fig. 5B**). Cluster 17 showed significant enrichment for the functional properties *sodium* (ion coupling), *SLC6* (family), *LeuT* (fold) and *amino acids, peptides and derivatives* (substrate class).

The annotated substrates of the SLCs in cluster 17 included different amino acids and derivatives (glycine for SLC6A5 and SLC6A9, GABA and betaine for SLC6A12, and neutral amino acids for SLC38A2, SLC38A4, SLC6A14, SLC6A15 and SLC6A19) as well as a sugar (myo-inositol for SLC5A3), while all of them co-transport Na^+^ (**Fig. 5B**). To pinpoint genes unique to this cluster, we examined the most strongly deregulated genes of cluster 17 and compared them to the average differential gene expression in other clusters (**Fig. 5C**). Many of the strongly and uniquely affected genes in this cluster encode SLCs: SLC5A3 (itself a cluster 17 member), SLC6A6, SLC16A6, SLC16A9, SLC16A14, SLC43A2 and SLC44A2. Hence, cluster 17 presented an example whereby the common metabolic effect of a group of SLCs led to a unified deregulation of another group of SLCs. Substrates are known for SLC5A3 (myo-inositol), SLC6A6 (taurine), SLC16A9 (carnitine), SLC43A2 (neutral amino acids) and SLC44A2 (choline) (Goldmann et al, accompanying manuscript). Although not demonstrated for human SLC16A6, the protein encoded by the mouse *Slc16a6* gene has been proposed to transport taurine (Higuchi *et al*, 2022). Meanwhile, our metabolomics data showed that overexpression of the orphan SLC16A14 reduced intracellular carnitine and propionylcarnitine levels (**Fig. 2D**), compatible with a role in carnitine efflux. Substrates of SLCs in cluster 17 and of the deregulated SLCs all have roles as osmolytes (Burg & Ferraris, 2008; Steeves & Baltz, 2005; Bussolati *et al*, 2001). We hypothesized that overexpression of the SLCs in cluster 17 commonly caused intracellular substrate accumulation, with intracellular osmolarity increased compared to the extracellular one, leading to hypoosmotic stress. The transcriptional response appeared to reflect cellular compensatory reactions and was consistent with a reduction of substrate osmolytes. This reduction was mediated by a series of changes: decrease intracellular taurine by downregulation of taurine importer SLC6A6 and upregulation of taurine exporter SLC16A6 as well as decreases in intracellular carnitine likely to be caused by upregulation of carnitine exporter SLC16A9 and the putative acylcarnitine exporter SLC16A14 (**Fig. 2D**). The likelihood of this interpretation is supported by the observed upregulation of the metabolic enzyme glycerophosphocholine phosphodiesterase 1 (GPCPD1), that converts glycerophophorylcholine (GPC) to choline. Upregulation of both GPCPD1 and SLC44A2 promotes GPC catabolism and choline export. Additionally, upregulation of SLC43A2, an amino acid uniporter, is expected to reduce excess intracellular amino acid levels.

To validate the metabolic changes in cluster 17 caused by SLC overexpression, we performed targeted metabolite measurements covering substrates of the deregulated gene products and substrates of the cluster members themselves, which were not measured in the original metabolomics panel. This was performed on five members of the cluster (SLC5A3, SLC6A5, SLC6A9, SLC38A2, SLC38A4) covering a diversity of SLC families and annotated substrates. We confirmed that known substrates of cluster members accumulated significantly upon doxycycline induction of the respective cell line: myo-inositol for SLC5A3, glycine for SLC6A5 and SLC6A9, and histidine as well as methionine for SLC38A2 and SLC38A4 (**Fig. 5D**). Substrates of the proteins encoded by the common deregulated genes were reduced too, aligning with a transcriptional program aimed at decreasing intracellular osmolyte levels: taurine (SLC6A6 and SLC16A6), carnitine and creatine (SLC16A9), and GPC (SLC44A2). Although regulation of transporter activity often resides on important translational and posttranslational mechanisms, these findings suggested that it was also possible to interpret changes in mRNA levels as part of metabolic regulatory circuits. In summary, SLCs in cluster 17 induced a transcriptional response to hypoosmotic stress, manifested in deregulation of SLCs and other genes associated with reduction in intracellular osmolyte levels (**Fig. 5E**). This highlighted how observed transcriptional changes corresponded to metabolite changes, supporting the validity of the transcriptomics data set in the broad coverage of metabolic pathways. The approach addresses properties in the regulation of metabolism that are complementary to those that can be inferred by our targeted metabolomics approach.

### A group of SLCs deregulating the cellular glycosylation machinery

Another transcriptomic cluster, cluster 6 was also significantly enriched in four different functional SLC properties: *SLC35* (family), *DMT* (fold), *Golgi* (localization) and *nucleosides, nucleotides and nucleotide-sugars* (substrate class) (**EV Fig. 5B**). Furthermore, the adjacent cluster 7 contained several SLC35 members and also showed significant enrichment for DMT fold and Golgi localization (**Fig. 6A**). Due to the high similarity in the enrichment of functional properties between these two adjacent clusters, we considered them as a larger cluster of SLCs with a common transcriptomic effect. The SLC35 family represents the major family of Endoplasmic Reticulum (ER)- and Golgi-localized nucleotide sugar transporters that govern the availability of substrates for glycosylation reactions and carry the DMT protein fold (Ferrada & Superti-Furga, 2022). In agreement with the enriched functional properties, gene set enrichment analysis (GSEA) on metabolic genes averaged across the cluster revealed GO Terms and Molecular Functions related to glycosylation, including glycosyltransferase and hexosyltransferase activity, as well as Golgi-associated GO Cellular Compartment terms (**Fig. 6B**). In line with these results, the metabolic profiles of SLCs from clusters 6 and 7 showed an increase in N-acetylglucosamine phosphate (GlcNAc-P), uridine monophosphate (UMP), cytidine monophosphate (CMP) and N-acetylneuraminic acid (Neu5Ac) (**Fig. 6C**), pointing to perturbations in nucleotide sugar synthesis and transport. GlcNAc-P and Neu5Ac belong to the connected metabolic pathways of hexosamine and sialic acid biosynthesis, which link to nucleotide metabolism through consumption of uridine triphosphate (UTP) and cytidine triphosphate (CTP) to produce nucleotide-sugars UDP-N-acetylglucosamine (UDP-GlcNAc), UDP-N-acetylgalactosamine (UDP-GalNAc) and CMP-N-acetylneuraminic acid (CMP-Neu5Ac), which are substrates for glycosylation reactions in the ER and Golgi (**EV Fig. 6A**). Multiple SLCs of the transcriptomic clusters 6 and 7 belong to the metabolomic cluster 5 (**EV Fig. 6B**, highlighted in black), indicating a high degree of interrelation between metabolic and transcriptional effects of SLCs. Taken together, the transcriptomic clusters 6 and 7 (thereafter referred to as the ‘glycosylation-related cluster’) displayed a common gene expression signature and altered metabolism related to altered cellular glycosylation.

**Figure 6.**
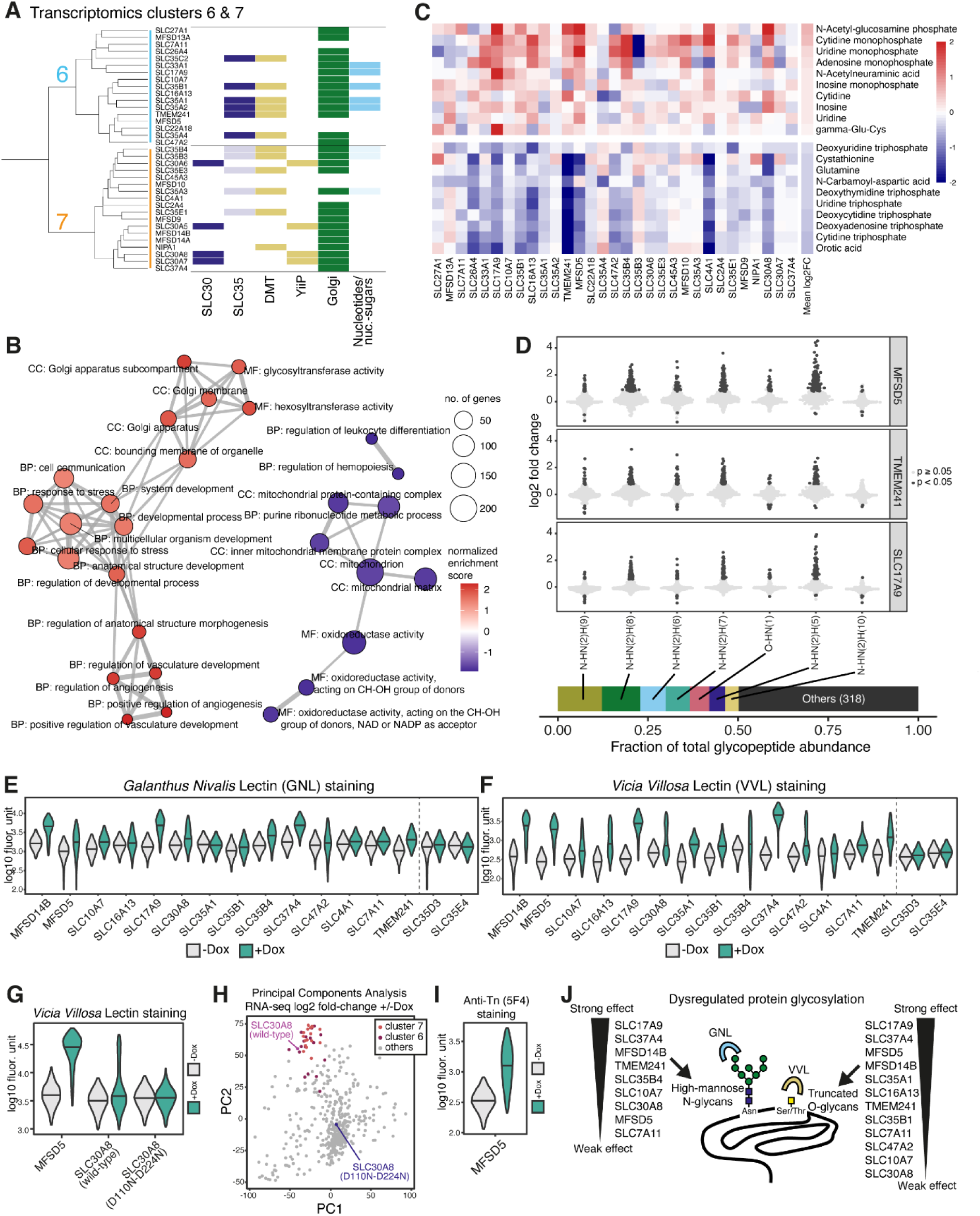
Multi-omics-based identification of an SLC cluster with effects on cellular N- and O-linked glycosylation. **(A)** Transcriptomic clusters 6 and 7 (thereafter referred to as combined ‘glycosylation-related cluster’) with significant SLC functional feature enrichment (Fisher’s test p < 0.2). **(B)** GO terms significantly enriched (p-adj. < 0.05) by gene set enrichment analysis (GSEA) on mean log2 fold-changes of metabolic genes across glycosylation-related cluster. **(C)** Heatmap of metabolomics data showing doxycycline (Dox) vs uninduced log2 fold changes for SLCs in glycosylation-related cluster. Only metabolites with 10 highest and lowest average log2FCs are shown. **(D)** Violin plots showing Dox-induced vs uninduced glycopeptide abundance log2 fold changes for 7 most abundant glycan compositions quantified by MS-based glycoproteomics. Individual glycopeptides are plotted as discrete points and colour coded by significance. Bottom: stacked barplot indicating fraction of total glycopeptide signal contributed by each glycan composition, showing that 7 most abundant glycan compositions account for 50% of the total glycopeptide abundance. **(E)** Intracellular GNL staining of +/- Dox cells following permeabilization, primary staining with biotinylated lectin and secondary staining with streptavidin-Alexa Flour 647. **(F)** Cell surface *Vicia Villosa* Lectin (VVL) staining of +/- Dox cells following primary staining with biotinylated lectin and secondary staining with streptavidin-Alexa Flour 647. **(G)** Comparison of cell surface VVL staining of MFSD5, SLC30A8 wild-type, and SLC30A8 D110N-D224N (transport deficient mutant) +/- Dox cells following primary staining with biotinylated lectin and secondary staining with streptavidin-Alexa Flour 488. **(H)** Principal components analysis of gene expression log2FC profiles of SLC30A8 D110N-D224N along with the 450 SLCs used in transcriptomic clustering. **(I)** Anti-Tn antibody staining of MFSD5 +/- Dox cells following primary staining with 5F4 and secondary staining with anti-mouse IgM Alexa Flour 568. For all flow cytometry violin plots, each plot represents flow cytometry measurements of at least 30 000 cells pooled from 3 replicate wells, and horizontal bisecting lines represent population medians. **(J)** Schematic illustration of effects on N- and O-linked glycosylation by members of glycosylation-related transcriptomic cluster as validated by lectin staining.

To investigate the specific glycosylation changes, we performed multiplexed mass spectrometry-based glycoproteomics on three selected members of the cluster: MFSD5, an orphan SLC; TMEM241, recently identified as a Golgi UDP-GlcNAc transporter; and SLC17A9, a vesicular ATP transporter (Zhao *et al*, 2023; Sawada *et al*, 2008). We identified 11,938 glycopeptides corresponding to 1,093 proteins. The most abundant glycoproteins were enriched for the GO terms “Cellular Compartments” associated with secretory and endolysosomal pathways, as well as the “Biological Processes” of vesicle-mediated transport, endocytosis and amide transport (**EV Fig. 6C**). Principal components analysis of glycopeptide abundance profiles clearly separated doxycycline-induced from uninduced samples, as well as TMEM241 from the other two SLCs (**EV Fig. 6D**).

We next examined changes in the most highly abundant glycan compositions, treating N- and O-linked glycans separately. Out of 325 glycan compositions, the top seven compositions representing 40% of the 11,938 unique glycopeptide species quantified in our data (**EV Fig. 6E**) accounted for more than 50% of the total glycopeptide signal (**Fig. 6D, bottom panel**). Six were N-linked HexNAc(2)Hex(5–10), likely representing high-mannose N-glycans with the structure GlcNAc(2)Man(5–10). The remaining O-linked HexNAc(1) likely represented GalNAc, that initiates mucin-type O-glycan synthesis. Respective glycan compositions were increased across MFSD5, TMEM241 and SLC17A9 WT-OE cell lines (**Fig. 6D**).

To validate the increase in high-mannose N-glycans, we performed flow cytometry upon intracellular staining with *Galanthus Nivalis* Lectin (GNL), which specifically binds terminal mannose on glycan structures (Bojar *et al*, 2022). We tested the SLCs found in both the transcriptomic glycosylation cluster and the metabolomic cluster 5 (**EV Fig. 6B, highlighted in black**), and additional selected cell lines from the transcriptomic cluster (**EV Fig. 6B, highlighted in grey**). SLC35D3 and SLC35E4, which are not part of the glycosylation-related cluster were included as Golgi-and ER-localized reference controls. We observed the largest increases of GNL staining upon Dox-induced overexpression SLC17A9, SLC37A4, MFSD14B, TMEM241, and SLC35B4 (Cliff’s delta > 0.5) (**Fig. 6E**, **Table 9**). On the other hand, the reference controls SLC35D3 and SLC35E4 showed relatively small effects (Cliff’s delta 0.17 and -0.11 respectively). We further investigated intracellular distribution of GNL-stained high-mannose proteins by immunofluorescence confocal microscopy. In uninduced MFSD5 WT-OE cells, GNL stained sharp punctate loci, which overlapped with the lysosomal marker LAMP1 indicating that GNL predominantly stained lysosome-targeted proteins. This was confirmed by colocalization of GNL staining with M6P-1, which detects the mannose-6-phosphate (M6P) modification that marks soluble proteins for lysosomal delivery (Müller-Loennies *et al*, 2010) (**EV Fig. 6F**). Upon doxycycline induction, GNL and M6P-1 staining became diffused suggesting that MFSD5 overexpression impacted M6P-dependent lysosomal targeting.

To validate the glycoproteomic result of increased O-linked GalNAc we used *Vicia Villosa* Lectin (VVL) which binds terminal GalNAc (Bojar *et al*, 2022). VVL staining was measured by flow cytometry upon overexpression of the SLCs of the transcriptomic glycosylation cluster. VVL staining increased substantially upon Dox induction (Cliff’s delta > 0.4) in all cell lines except for SLC4A1 (Cliff’s delta 0.18), SLC35B4 (Cliff’s delta 0.16) and the reference controls SLC35D3 and SLC35E4 (Cliff’s delta 0.13 and 0.07 respectively) (**Fig. 6F**, **Table 10**). Closer examination of SLC47A2 and SLC30A8 revealed bimodal distributions in VVL staining signal and the presence of subpopulations with substantially elevated signal (**EV Fig. 6G**). This may suggest that these two SLCs deregulate glycosylation in a stochastic manner affecting only a subset of cells. In a transport-deficient mutant of SLC30A8 (Azzollini *et al*, 2024), the subpopulation with increased VVL staining was absent (**Fig. 6G, EV Fig. 6H**). In line with this result, the transcriptional profile of the SLC30A8 mutant was distinct from the SLCs of the glycosylation-related cluster (**Fig. 6H**), providing evidence that the transcriptional and glycosylation effects of SLC30A8 are dependent on transport functionality. O-linked GalNAc is also known as Tn antigen, a recognized tumour marker (Chia *et al*, 2016). We confirmed that cell surface binding of an anti-Tn antibody (5F4) increased upon doxycycline induction of MFSD5 WT-OE cells (**Fig. 6I**).

In summary, our family-wide omics data enabled the assignment and subsequent validation of a glycosylation-related function to the orphan SLC MFSD5 using the guilt-by-association principle. We identified a group of SLCs with diverse annotated functions but sharing an altered metabolite and gene expression signature, either stemming from or leading to an impact on the glycosylation machinery (**Fig. 6J**). Their commonality in driving increased O-linked GalNAc and N-linked high-mannose glycans offers avenues for future studies and mechanistic exploration.

## Discussion

By size, implication in human disease, and potential pharmacological exploitation, the target class of solute carriers (SLCs) compares well to GPCRs, kinases, proteases and ion channels (Cesar-Razquin *et al*, 2015; Oprea *et al*, 2018). Most of the functional studies that focus on a whole target class using CRISPR/Cas9 or siRNA use one specific readout or a few readouts in parallel (DelRosso *et al*, 2023; Timms *et al*, 2023). Less frequently, unbiased functional readouts with hundreds of genes of related function or structure are employed (Liu *et al*, 2023; Johnson *et al*, 2023). This work provides a first, although coarse, functional annotation by multi-omics of the entire SLC superfamily under comparable conditions. The parallel evaluation of changes in metabolites and transcripts across hundreds of isogenic cell lines each bearing altered expression of only one gene, offers data for hundreds of SLC transporters for which no functional annotation has been available.

This study represents a systematic survey of metabolic consequences of SLC function across the entire SLC superfamily, enhancing our understanding of the link between SLC activity and metabolism. We can consider targeted metabolomics as a quantitative ‘fingerprinting’ approach, that enables profiling and comparison at such a large scale. At the same time, metabolite coverage is limited and obviously fails to capture the full diversity of substrates transported by the SLC superfamily. Additionally, detection of a substrate is affected by multiple biological factors, such as transport kinetics, expression of the SLC, functional redundancy, metabolic conversion of the substrate, and metabolic need, which itself depends on cell type and cellular state. Despite these limitations, we found changes in annotated substrates or direct metabolic conversions for 102 SLCs (**Fig. 1E, F**; **Table 3**). Many of the remaining dysregulated metabolite hits might represent unannotated true substrates and we anticipate that our study will accompany follow-up validation efforts that identify novel substrates for both orphan and well-studied SLCs. Metabolic fingerprinting as performed here provides an overview of the metabolic effects caused by SLC expression and insight on the cellular control of metabolite levels. The magnitude of changes across more than 370 cell lines can indicate a window of tolerance for individual metabolite fluctuations that might be tighter or looser (**EV Fig. 2A**), possibly connecting to a metabolite’s regulatory role in feedback or feedforward mechanisms. The dataset offers the unique possibility to study correlations in metabolite abundance upon hundreds of perturbances in isogenic settings, perhaps leading to new insights into the plasticity of cellular metabolism in human cells.

For our large collection of SLC perturbation cell lines we chose SLC knock-out clones over knock-down pools to directly attribute observations to gene loss and ensure consistency across multiple measurements and assays. Using multiple clones for each knock-out cell line and considering clonal effects within the statistical analysis of metabolomics data mitigated clonal variance. Overexpression models allowed rapid SLC expression changes in a loss-of-function background, minimizing the kind of metabolic adaptation and possible genetic drift associated with the generation of cell lines. Nonetheless, protein overexpression can present serious drawbacks such as overload of the cellular machinery and functional consequences unrelated to physiological function. Some of the transcriptional programs we observe are likely elicited by protein stress.

Selecting the cell line models was a major decision at consortium level, aiming for a balance between physiological relevance, i.e. endogenous expression of the individual SLC gene, technical feasibility regarding the generation of cell lines, suitability for a commensurate data acquisition throughput, time, costs and robustness across experimental locations. We selected the colon cancer cell line HCT116 as our primary cell model because of its biological relevance in the transporter field and technical feasibility for large scale genome engineering. For SLCs that are not expressed in HCT116 or only at very low levels, compared to the other cancer cell lines in our panel, we chose a different cell model accordingly to provide a biologically more relevant model system (**EV Fig. 1A, 1B**). Additionally, we selected the Jump-In™ T-REx™ HEK293 cell line model to create a collection of overexpression cell lines for the entire SLC superfamily in an isogenic background. Nearly a quarter of the SLC superfamily were not studied in their natural cellular context, potentially affecting their functionality. However, we found literature-annotated substrates independently of whether an SLC was endogenously expressed in the given cell line. Using different cell line models each with their own standard culture conditions further challenged the complex metabolomics data analysis. A further constraint of our study was the necessity to limit to only standard culture conditions due to the already large sample number, fully aware that a unified standard media would unlikely represent optimal transport conditions for the different SLCs (Cantor *et al*, 2017). Without any doubt, this study represents but a first-pass survey, and more dedicated studies in more physiological settings will unravel subtleties and specificities not accessible in the present large-scale set-up.

Investigating functional profiles of individual SLCs revealed a novel role for SLC45A4 in mediating cellular export of polyamines, including putrescine and spermidine. So far, there is no annotated substrate for human SLC45A4, and evidence is scarce for the other three members of the SLC45 family. The heterologous expression of mouse *Slc45a2-4* increased sucrose uptake in yeast cells (Bartölke *et al*, 2014). Since we did not observe increased sucrose uptake in SLC45A4 overexpressing cells (**EV Fig. 3C**), human SLC45A4 may have an alternate function compared to its mouse homologue. Rat SLC45A1 was suggested to mediate transport of glucose and galactose, but not sucrose in mammalian cells (Shimokawa *et al*, 2002). The discrepancy between sugars transported by SLC45A1 and SLC45A2-4 in the different models may be due to differences in substrate specificity, but our results also raise the possibility that sugar transport may be a secondary effect of polyamine transport. Further biochemical binding assays using recombinant protein could further explore polyamine transport by SLC45A4, as will the three-dimensional structure.

For the identification of functional relationships between SLCs, we tested several strategies with varied sample normalization, clustering method and gene/metabolite sets, using the mean cluster stability and enrichment of functional SLC properties for evaluation. A limiting factor in the definition of functional clusters by metabolic profiles was the sparseness of the data set (142 metabolites, 378 SLCs) as well as the relatively high level of noise, as no threshold for significance was set. We did not face these limitations in the clustering of transcriptional profiles and therefore used transcriptomic clusters as starting points for investigations and metabolomic analyses of the respective groups of SLCs for complementation and refinement. This underscores the importance of considering complementary sets of information to identify and validate clustering patterns effectively.

From transcriptional profiling of 441 SLCs, we identified cluster 17 of Na^+^/osmolyte symporters with a shared deregulation of osmolyte transporters, likely aimed at balancing intracellular osmolyte levels and relieving hypoosmotic stress. We speculate that Na^+^ is the driving force in the accumulation of osmolytes by these Na^+^-dependent cotransporters, since DMEM (Sigma) contains a high concentration of Na^+^ ions (154 mM total Na^+^ from sodium chloride and sodium bicarbonate). The intriguing question arises as to why other Na^+^-dependent transporters of potential osmolytes, such as SLC6A6 (taurine), SLC6A8 (creatine), and SLC22A5 (carnitine), were not found in the same cluster. Among many possible explanations, we suspect that differences in substrate levels in the medium may be the cause: base DMEM contains 400 µM glycine, 200 µM of both histidine and methionine, and 40 µM myo-inositol, but no taurine, creatine or carnitine. Targeted metabolomics analysis of our culture media showed low levels of taurine (< 10 µM) likely from fetal bovine serum, while creatine and carnitine were below the limit of detection (**Table 11**). Therefore, substrate levels may be too low for substantial intracellular accumulation and subsequent induction of osmotic stress response upon SLC6A6, SLC6A8, or SLC22A5 overexpression. Among the commonly deregulated genes of cluster 17, taurine transporter SLC6A6 and myo-inositol transporter SLC5A3 are regulatory targets of the tonicity-responsive enhancer binding protein (TonEBP), a transcriptional regulator also known as NFAT5 (Burg & Ferraris, 2008). Regulatory mechanisms have not been reported for the other deregulated genes SLC43A2, SLC16A6, SLC44A2, SLC16A14, SLC16A9. We expect that unravelling the common regulatory mechanism is likely to shed light on cellular responses to hypoosmotic stress and represents an intriguing subject of future investigation. In addition, considering the constraints due to media composition, such a regulatory mechanism may be affecting yet more SLCs.

Further we identified an apparently diverse group of SLCs, including the UDP-GlcNAc transporter TMEM241, vesicular ATP transporter SLC17A9, zinc transporter SLC30A8, several members of the SLC35 family and the orphan MFSD5, with a shared effect on dysregulating the cellular glycosylation machinery. It is not surprising that overexpression of SLC35 family members dysregulate glycosylation, given their central role in transport of nucleotide sugars into the ER and Golgi for glycosyltransferase reactions (Hadley *et al*, 2019). Less recognized are the links between vesicular ATP transport (SLC17A9) or zinc transport by SLC30 family members, of which four are present in the cluster (SLC30A5–8). The SLC30 family is primarily involved in the transport of zinc out of the cytoplasm, either into extracellular space or intracellular compartments (Chen *et al*, 2024). Overexpression of SLC30 family members possibly leads to Zn^2+^ accumulation in the secretory pathway which may perturb glycosylation (Rømer *et al*, 2023). Given the important role of SLC17A9-mediated ATP transport in the secretory and endolysosomal pathways, both closely connected to the Golgi, where glycan biosynthesis and breakdown takes place, SLC17A9 overexpression may precisely impact the crosstalk between these cellular compartments. The orphan transporter MFSD5 shows high structural similarity to SLC17A9 (Ferrada & Superti-Furga, 2022) and was reported to bind ATP (Jelcic *et al*, 2020), while plant homologs of this SLC are reported as Golgi-localized transporters of S-adenosylmethionine (Temple *et al*, 2022). Based on these observations, we speculate that MFSD5 may mediate transport of ATP or other adenosyl compounds, although this hypothesis remains to be tested.

Common glycosylation changes by this group of SLCs include elevated high-mannose N-glycans and Tn antigen levels. Increased high-mannose N-glycans are frequently upregulated in cancer and correlate with cancer grade (Chatterjee *et al*, 2021), while the Tn antigen is a well-recognized oncofetal antigen expressed in a large proportion of solid tumours including 70-90% of breast colon, lung, bladder, cervix, ovary, stomach, pancreatic, and prostate cancers (Ju *et al*, 2011; Chia *et al*, 2016; Mereiter *et al*, 2016) and supports tumour progression through immunomodulatory interactions (van Vliet *et al*, 2006). The mechanisms driving these cancer-associated glycosylation changes are not fully understood but hold potential for therapeutic intervention (Chia *et al*, 2016; Mereiter *et al*, 2019). Mechanistic elucidation of the SLC-driven glycosylation changes discovered in this study is expected to provide new insights into the roles of transporters in glycosylation and may reveal new biomarkers or targets for cancer therapy.

The analyses conducted serve as exemplary explorations of this rich resource, providing a foundation for numerous future studies. Given the significance of metabolism in physiology and disease, understanding the roles of individual metabolites, their regulation, and their function as regulators in an integrated manner is paramount. SLCs are attractive pivotal players to modulate metabolism as they often represent accessible “gates” and are eminently druggable (Lin *et al*, 2015; Wang *et al*, 2020; Dvorak & Superti-Furga, 2023).

The combination of metabolomic and transcriptomic data sets holds considerable promise for future research, once the connections between metabolism, transcription, and expression are better understood. We anticipate that these data sets will serve as valuable resources for understanding metabolic conditions in disease contexts and the compensatory interactions influencing disease. This work together with the accompanying studies on the SLC knowledgebase (Goldmann *et al*), the SLC genetic interactions (Wolf and Leippe *et al*) and the SLC protein-protein interactions (Frommelt and Ladurner *et al*) is a series of four publications which presents the largest study on the SLC superfamily to date and possibly a unique case of concerted functional assignment to a large superfamily of human genes. The underlying reagents and data sets are publicly accessible through a web portal and public repositories (https://re-solute.eu/; https://www.addgene.org/depositor-collections/re-solute/; https://www.atcc.org/cell-products/cell-models/solute-transporter-carrier-cells).

## Materials and Methods

### Cell line generation

HCT116 (RRID:CVCL_0291) and LS180 (RRID:CVCL_0397) cell lines were cultured in RPMI (R8758 Sigma) + 10% Fetal Bovine Serum (10270-106, Lot 42F8381K, Gibco) + Penicillin-Streptomycin (15140-122, Gibco). Jump-In™ T-REx™ HEK293 (RRID:CVCL_YL74), HuH-7 (RRID:CVCL_0336), 1321N1 (RRID:CVCL_0110) and SK-MEL-28 (RRID:CVCL_0526) in DMEM (D5796 Sigma) + 10% Fetal Bovine Serum (10270-106, Lot 42F8381K, Gibco) + Penicillin-Streptomycin (15140-122, Gibco). All cell lines were grown at 37 °C and 5% CO2. Transgene expression is induced via doxycycline treatment (1 µg/ml (D9891-1G, Sigma Aldrich; CHEBI:50845).

To generate monoclonal KO clones, cell lines were stably transduced with the lentiviral vector pLentiCRISPRv2 (Addgene #52961) expressing gRNAs targeting SLCs (**Table 1**). After 5-10 days puromycin selection, cell pools were sorted to single cells using FACS or a limited dilution series and monoclonal cell lines were expanded and genotyped using an NGS-based approach (Veeranagouda *et al*, 2018). At least two independent KO clones were generated for each target. A detailed description of the KO generation can be found here: https://zenodo.org/records/7457297.

To re-express functional and tagged SLCs in the respective KO background, codon-optimized SLC cDNAs were cloned from pDONR221 gateway entry vectors (https://www.addgene.org/depositor-collections/re-solute/) into doxycycline-inducible lentiviral gateway destination vectors containing C- or N-terminal HA-Twin-Strep® tags (Addgene # 194066, 194065) and stably transduced into KO cell lines. For N-terminally tagged SLCs, cDNAs with STOP codons at the end of the SLC ORF were used. Transgene expression was induced via doxycycline treatment at 1 µg/ml (D9891-1G, Sigma Aldrich; CHEBI:50845) for 24 hours. For each SLC, two pooled KO-OE cell lines were generated, each derived from an independent clonal KO cell line. A more detailed description of the KO-OE cell line generation can be found here: https://zenodo.org/records/7457295.

To generate cell lines overexpressing codon-optimized tagged SLCs in a WT background (WT-OE), SLC cDNAs were cloned from the same pDONR221 vectors into modified pJTI™ R4 CMV-TO MCS pA vectors (Thermo Fisher Scientific) that contained C- or N-terminal HA-Twin-Strep® tags. Jump-In™ T-REx™ HEK293 cells (Thermo Fisher Scientific) were stably transfected with these constructs as recommended by the manufacturers protocol. Transgene expression was induced via doxycycline treatment at 1 µg/ml for 24 hours or the duration indicated. A more detailed description of the pooled WT-OE cell line generation and quality control can be found here: https://zenodo.org/records/7457221 and https://zenodo.org/records/5566805.

### Cell line selection for this study

For several SLC, multiple cell lines and transcriptomics data sets were generated. Based on expression analysis by western blot, immunofluorescence and AP-MS proteomics, we chose one cell line for each SLC to be included in this study. For six SLCs we were not able to generate and include a functional cell line. For metabolite extraction, the appropriate cell line model for each SLC was chosen based on its expression across the panel of cell lines described above. In case the SLC is endogenously expressed in one of the six cancer cell lines and a respective knock-out cell line was available, the KO-OE cell line model was chosen for targeted metabolomics analysis. In case the SLC is not endogenously expressed in any of the cancer cell lines, or it was not possible to derive a respective knock-out cell line, the WT-OE model was chosen.

### Metabolite extraction

Cells were plated in quadruplicates in 6-well or 24-well plates in culture medium in the absence or presence of 1 μg/ml doxycycline (150 000 cells per well for HCT 116 and LS180, 100 000 cells per well for 1321N1, 50 000 cells per well for Huh-7, 70 000 cells per well for SK-MEL-28, 750 000 / 150 000 cells per well in poly-L-lysine coated 6-well / 24-well plates for Jump-In™ T-REx™ HEK293). After 16- 24 h, cells were first gently washed with room temperature ammonium carbonate buffer (75 mM NH4HCO3, pH 7.4). Then, cells were transferred to ice, where 300 µl/well in 24-well plate or 1500 µl/well in 6-well plate of ice-cold 80:20 MeOH:H2O was added. The cells were then scraped and transferred to a pre-cooled Eppendorf tube and immediately snap frozen in liquid nitrogen. Samples were thawed on ice before being centrifuged at 16 000 × g for 10 min at 4 °C. The clarified metabolite-containing supernatants were moved into a high-performance liquid chromatography vial and stored at -80 °C until further preparation for LC-MS/MS analysis. Cell extracts were dried using a nitrogen evaporator. The dry residue was reconstituted in 16 µl H2O and 4 µl of sample extract was used for LC-MS/MS analysis. A mixture of isotopically labelled internal standards (Synthetic heavy isotope labelled metabolites from Sigma-Aldrich and Cambridge Isotope Laboratories as well as Metabolite Yeast Extract (U-13C, 98%) from ISOtopic Solutions, Vienna, Austria) was added to each sample either with the extraction solvent or with the H2O at reconstitution.

### Targeted metabolomics

A 1290 Infinity II UHPLC system (Agilent Technologies) was used for the chromatographic separation utilizing a ZORBAX RRHD Extend-C18, 2.1 × 150 mm, 1.8 µm analytical column (Agilent Technologies) and a SecurityGuard ULTRA Cartridge UHPLC C18 2.1 mm ID precolumn (Phenomenex). The columns were maintained at a temperature of 40 °C. The mobile phase A was 3% MeOH (v/v), 10 mM tributylamine, 15 mM acetic acid in H2O, and mobile phase B was 10 mM tributylamine, 15 mM acetic acid in MeOH. The gradient elution with a flow rate 0.25 mL/min was performed for a total time of 24 min followed by back flushing of the column using a 6 port 2-position divert valve for 8 min using acetonitrile. At the end of the run 8 min of column equilibration with 100% mobile phase A was performed. Chromatographic separation was coupled to a 6470 triple quadrupole mass spectrometer (Agilent Technologies). The triple quadrupole mass spectrometer was operated in electrospray ionization negative mode, spray voltage 2 kV, gas temperature 150 °C, gas flow 1.3 L/min, nebulizer 45 psi, sheath gas temperature 325 °C, and sheath gas flow 12 L/min. The metabolites of interest were detected using a dynamic Multiple Reaction Monitoring (dMRM) mode.

### RNA isolation

For RNA isolation 2.5 × 10^5^ cells per well were seeded in four wells of a 12-well plate. In two of these wells, complete growth media is added, in the other two wells complete media containing 1 µg/ml doxycycline (Sigma Aldrich; CHEBI:50845). Cell lines were harvested 24 hours after doxycycline induction. At the time of harvest, media is removed from cells and cells are washed once with 1 ml PBS buffer (D8537, Sigma). After washing, 200 µl buffer RLT (Qiagen) is added directly to the cells. After complete cell lysis, lysates are transferred into matrix tubes and frozen at -80 °C until further treatment. For RNA purification, RNeasy 96 Kits (Qiagen) were used. DNA is removed by on-column DNase treatment as described in the manual. After purification, RNA is eluted in 60-80 µl RNase-free water and stored at -80 °C until further usage. RNA concentrations are measured using the QuantiFluor® RNA System (Promega). For a selection of samples, RNA integrity was determined on a Bioanalyzer (Agilent). Total RNA samples were quantitatively and qualitatively assessed using the fluorescence-based Broad Range Quant-iT RNA Assay Kit (Thermo Fisher Scientific) and the Standard Sensitivity RNA Analysis DNF-471 Kit on a 96-channel Fragment Analyzer (Agilent), respectively.

### Library preparation and RNA sequencing

RNA samples were normalized on the MicroLab STAR automated liquid platform (Hamilton). Total RNA input of 250 ng was used for library construction with the NEBNext Ultra II Directional RNA Library Prep Kit for Illumina #E7760, together with the NEBNext Poly(A) mRNA Magnetic Isolation Module #E7490 upstream and the NEBNext Multiplex Oligos for Illumina #E7600 downstream (all New England Biolabs). The only deviation from the manufacturer’s protocol was the use of Ampure XP beads (Beckman Coulter) for double-stranded cDNA purification, instead of the recommended SPRIselect Beads. The index PCR was performed with 12 cycles, while the final library was eluted in 35 µL. Library preparation was performed either manually or automated using a Biomek i7 Hybrid workstation (Beckman Coulter) as described previously (Santacruz *et al*, 2022). mRNA libraries were then quantified by the High Sensitivity dsDNA Quanti-iT Assay Kit (ThermoFisher) on a Synergy HTX (BioTek). mRNA libraries were also assessed for size distribution and adapter dimer presence (<0.5%) by the High Sensitivity Small Fragment DNF-477 Kit on a 96-channel Fragment Analyzer (Agilent).

All sequencing libraries were normalized on the MicroLab STAR (Hamilton), pooled and spiked in with PhiX Control v3 (Illumina). The library pools were subsequently clustered on S4 Flow Cell and sequenced on a NovaSeq 6000 Sequencing System (Illumina) with dual index, paired-end reads at 2 x 50 bp length (Read parameters: Rd1: 51, Rd2: 8, Rd3: 8, Rd4: 51), aiming for an average depth of 25 million Pass-Filter reads per sample.

### Metabolomics data processing and differential analysis

PeakBotMRM (https://github.com/christophuv/PeakBotMRM) was utilized for processing the raw LC-MS/MS data, followed by manual correction of some incorrectly recognized peaks and peak boundaries using the graphical user interface.

Metabolite peak areas were normalized to those of reference internal standards. Forty-seven metabolites had internal standards with the same chemical structures, and therefore these respective internal standards were used as references. For the remaining metabolites we performed weighted linear regression on the expected metabolite concentrations against the calculated ratios for each metabolite and each internal standard. When internal standards achieved an adjusted R2 greater or equal to 0.85 in at least 75% of the runs in which the metabolite was measured, the internal standard with the highest percentage was selected as the reference. If multiple internal standards achieved the same highest percentage, all were selected. For ten metabolites an adjusted R2 greater or equal to 0.85 was achieved in less than 75% and again the internal standard with the highest percentage was chosen. Individual injections were removed, or a reduced set of internal standards was used for normalization in case certain internal standards were not detected in certain runs. Normalization was performed by calculating the ratio of a metabolite peak area to the corresponding internal standard peak area, and subsequent log-transformation. For metabolites with multiple reference internal standards, the results were averaged. To also enable comparisons between different runs, we corrected for run-to-run differences (batch effects) by centering and scaling the internal-standard normalized values of each metabolite, per run, to its overall (i.e. across runs) median and interquartile range, respectively.

Finally, differential abundance upon SLC overexpression was tested for each metabolite by contrasting its normalized and batch-corrected values in doxycycline induced samples to the corresponding uninduced samples in a one-way ANOVA for WT-OE cell lines, or a two-way ANOVA for KO-OE cell lines, modeling the clone effect as additional factor. Resulting p-values were corrected for multiple testing per SLC, using the Benjamini-Hochberg procedure (Benjamini & Hochberg, 1995). Differentially abundant metabolites were called at a false discovery rate of 5%.

### Transcriptomics data processing and differential analysis

RNA-Seq data was processed using a Snakemake (version 6.6.1) pipeline running on Python (version 3.7). The pipeline comprised i) the read trimming using cutadapt (version 2.8), ii) the mapping of paired-end reads to the reference genome (GRCh38.p13 assembly with Ensembl 98 annotation) and to the overexpressed codon-optimized cDNA sequences for 447 SLCs using STAR (version 2.7.9a), iii) the quantification of reads per gene model using RSEM (version 1.3.2) as well as iv) the quality control of the sequencing data using FastQC (version 0.11.9) and the quality control before and after alignment using RSeQC (version 4.0.0) ensuring reliable downstream analyses. Quality control results after the alignment and quantification were compiled to individual reports using MultiQC (version 1.11).

### Differential expression analysis

To identify changes in gene expression levels upon overexpression of an SLC, differential expression analysis was performed by contrasting gene read counts between doxycycline-induced versus uninduced samples of the same cell line (2 replicates each). Genes with a total of less than 10 reads in all analyses were excluded as well as the codon-optimized cDNA sequences for 447 SLCs to consider only endogenously expressed SLC sequences. The DESeq2 package (version 1.42.1; (Love *et al*, 2014)) in R (version 4.3.3) was used to build a gene-level read count model, where counts were normalized by sample-specific size factors to account for differences in library size. Estimated gene-wide dispersions were used for the count modeling and a Generalized Linear Model was fit for each gene followed by hypothesis testing for differential expression using the Wald test. Resulting p-values were corrected for multiple testing using the Benjamini-Hochberg procedure (Benjamini & Hochberg, 1995) and shrinkage of log fold changes was performed using the apeglm method (Zhu *et al*, 2019). Differentially expressed genes were called at a false discovery rate of 5%.

### Uniquely deregulated genes

To find genes that are strongly differentially regulated in specific cell lines, we calculated z-scores of the shrunken log fold change of each gene within a certain analysis and also across all analyses. Genes were then further filtered for significance of differential expression (adjusted p-value < 0.05) and for minimal signal (the gene had to have in one condition at least 50 read counts in both replica).

### Reactome pathway analyses

Pathways, reactions, compounds and proteins were extracted from the human Reactome pathway database (version 87), downloaded in BioPAX format. The data set was limited to “Metabolism” pathway (R-HSA-1430728) and its subcomponents.

Metabolites targeted in the metabolomics method were matched to compounds annotated in Reactome if they represented a more generic or a more specific term in the ChEBI ontology (release 229), considering the following relations: “is_a”, “is_conjugate_base_of”, “is_conjugate_acid_of”, “is_tautomer_of”, “is_enantiomer_of” and “has_role”. Genes measured in transcriptomics analyses were matched to proteins annotated in Reactome via HGNC database (downloaded 2024-01-17).

To assess the regulation of individual metabolic pathways, we counted for each pathway the number of transcriptomics and metabolomics analyses featuring differentially regulated genes or metabolites that were matched to any reaction within the pathway. As pathways higher up in hierarchy or pathways using more generic compounds will be found regulated very often, we tested for significance of the regulation frequency by permutation test, shuffling the identities of the quantified metabolites and genes 200 000 times. The resulting significance levels were visualized on a Voronoi treemap of the hierarchical structure of all sub-pathways of human Metabolism pathway in Reactome, to show areas specifically affected by SLC overexpression.

To check whether we find known SLC substrates regulated in the targeted metabolomics experiments, we matched the targeted metabolites to the annotated substrates of SLCs (Goldmann et al, accompanying manuscript), using the ChEBI ontology in the same way as described above.

Extending this approach to also consider metabolic conversion products of the annotated substrates, we searched for Reactome reactions that featured an annotated substrate (or a more specific term) matched to a reaction educt, and a targeted metabolite (or a more specific term) matched to a reaction product. Matching was done via ChEBI ontology using the same relations as listed above. Educts that were also products of the same reaction, as well as the substrate annotations of “hydron”, “hydroxide” and “water” were not considered.

### Hierarchical clustering

Clustering was performed on 381 metabolic and 450 transcriptional profiles of metabolites and genes which were measured in all analyses (142 metabolites, 14,187 genes). The profiles of shrunken log fold changes were standard normalized, and Euclidean distance between the profiles was used for a hierarchical clustering (hclust function in R, version 4.3.3, with Ward’s method).

As a measure of functionality, we tested clusters for enrichment of SLC functional properties of different classes (coupled ion, family, fold, location, substrate class). Respective classes and annotations are described in Goldmann et al, accompanying manuscript. Fisher’s exact test for overrepresentation was performed for each functional property in each cluster, only considering properties with at least 3 annotations. Resulting p-values were corrected for multiple testing using the Benjamini-Hochberg procedure (Benjamini & Hochberg, 1995) for each class separately, and enrichments of functional SLC properties in clusters were called at a false discovery rate of 20%.

Guided by the mean silhouette width (cluster library, version 2.1.6; (Rousseeuw, 1987)) and the enrichment analysis, the metabolomics and transcriptomics dendrograms were divided into 48 and 60 clusters respectively.

### Transcriptomics-based assessment of metabolic pathway deregulation

To assess changes to expression of metabolic genes driven by each SLC and transcriptomic cluster, we mapped genes to Reactome pathways as described above. For each cell line, we then calculated the mean expression log2 fold change of all associated genes belonging to each pathway. We defined top-level pathways as the set of direct child pathways of the human Metabolism pathway (R-HSA- 1430728), and selected the 9 pathways with largest number of associated genes for plotting on **Fig. 5A**.

### GO term enrichment analysis

To determine enriched GO terms within the overall gene expression profile of transcriptomic clusters 6 and 7, Gene Set Enrichment Analysis (GSEA) was performed in R using the *clusterProfiler* package (Yu *et al*, 2012). Standard-normalized log2FCs were averaged across SLCs of the two clusters, arranged in descending order and pre-ranked GSEA performed with the *gseGO* function using the following parameters: ont = “ALL”, keyType = “ENSEMBL”, minGSSize = 10, maxGSSize = 500, OrgDb = “org.Hs.eg.db”, by= “fgsea”.

### Targeted LC-MS/MS for metabolite quantification

For stable isotope labeling experiment, HCT116 SLC45A4 KO-OE cells were grown in RPMI media supplemented with either 4 mM ^13^C^15^N-arginine or ^13^C^15^N-glutamic acid for 3 days, then induced with 1 µg/mL Dox for 24 h. For other experiments, SLC45A4 KO-OE cells were seeded in regular RPMI media overnight, then switched to RPMI containing 10% regular FBS, dialyzed FBS, horse serum or no serum, with supplementation of 1 mM DFMO, 2 mM aminoguanidine, or 0.1 unit/mL porcine diamine oxidase (Sigma), and +/- 1 µg/mL Dox for 9 h. Metabolite extraction was performed as described above, and additionally the protein pellets remaining after extraction were dissolved in 0.5 M KOH and subsequently quantified by BCA assay.

Targeted metabolite measurements were performed on a Waters Xevo TQ system utilizing an ACQUITY UPLC® HSS T3 1.8µm reversed-phase column (Waters) maintained at 40 °C. The mobile phase A was water with 0.1% formic acid, and mobile phase B was acetonitrile with 0.1% formic acid. The elution gradient was as follows: t = 0, 2% solvent B, flow rate 0.15 mL/min; t = 2.5, 2% solvent B, flow rate 0.15 mL/min; t = 3.5, 100% solvent B, flow rate 0.3 mL/min; t = 6, 100% solvent B, flow rate 0.3 mL/min; t = 6.1, 2% solvent B, flow rate 0.2 mL/min; t = 8, 2% solvent B, flow rate 0.15 mL/min. The triple quadrupole mass spectrometer was operated in electrospray ionization mode with polarity determined according to the analyte. Other settings were as follows: capillary voltage 3 kV, source temperature 150 °C, desolvation temperature 350 °C, desolvation gas flow 650 L/h, collision gas flow 0.15 mL/min. MRM transitions were determined for each target metabolite using the IntelliStart function of MassLynx software and were as follows: GABA (+) 104.0953 → 69.0814; putrescine (+) 89.0681 → 72.06; spermidine (+) 146.2537 → 72.1248; ^13^C^15^N-Glutamic acid (+) 154.1596 → 89.1233; sucrose (-) 341.1481 → 179.0127. Waters RAW data folders were converted to mzML files using Proteowizard MSConvert (Chambers *et al*, 2012) and peak areas were quantified using EL-MAVEN (Agrawal *et al*, 2019). With exception of the stable isotope labeling experiment, peak areas were then normalized to both internal standard (^13^C^15^N-glutamic acid) and BCA protein quantification values and the results expressed as normalized peak areas.

### SLC45A4 expression, purification, proteoliposome preparation, and putrescine uptake assay

SLC45A4 was expressed and purified as previously described (Raturi *et al*, 2023). Briefly, full-length human SLC45A4 was expressed in Expi293F cells using the BacMam system. Cells were harvested after 48 h and resuspended in lysis buffer (300 mM NaCl, 50 mM HEPES pH 7.5) in the presence of cOmplete Protease Inhibitor Cocktail tablets (Roche) and lysed at 4 °C by two passes through an EmulsiFlex-C5 homogenizer (Avestin). The lysate was clarified by centrifugation at 10 000 × g for 15 min, and membrane pelleted by ultracentrifugation at 100 000 × g for 1h at 4 °C. The resulting membrane pellet was solubilized in lysis buffer supplemented with 1% dodecylmaltoside (DDM) and 0.1% cholesterol hemisuccinate (CHS). The solubilized membranes were then incubated with pre-equilibrated Strep-Tactin XT Superflow resin (IBA-Lifesciences) for 1h at 4 °C, washed with column buffer (300 mM NaCl, 50 mM HEPES pH 7.5, and 0.05% DDM/CHS (Anatrace)) supplemented with 10 mM MgCl2 and 1 mM ATP, and protein was eluted with column buffer supplemented with 50 mM D-biotin. The affinity tag was cleaved with TEV protease overnight and the protein further purified by reverse IMAC purification using TALON resin (Takara). The tag-cleaved protein was concentrated using a 100 kDa cutoff centrifugal concentrator (Sartorius) and subjected to size exclusion chromatography using a Superdex 200 10/300 GL column (GE Healthcare) pre-equilibrated with gel filtration buffer (150 mM NaCl, 20 mM HEPES pH 7.5, 0.025% DDM/CHS). Peak fractions were pooled for subsequent experiments.

Liposomes were prepared by dissolving 120 mg soy lipids (Sigma-Aldrich) in 20 mL chloroform and then drying as a thin film in a rotovap. The lipids were resuspended in 6 mL of encapsulation buffer (20 mM HEPES pH 7.5, 150 mM NaCl, 5 mM glycine and 5 mM valine), then subjected to 10 freeze-thaw cycles by transferring between liquid nitrogen and 37 °C water bath. To form liposomes of uniform size, lipids were extruded 21 times through a 400 nm filter membrane using an LF-1 handheld syringe extruder (Avestin). Triton X-100 was added to a final concentration of 0.1%, then purified SLC45A4 protein added at a ratio of 0.1 mg protein per 4 mg lipid. For the control liposomes, an equivalent volume of encapsulation buffer was added to the liposomes. The mixture was incubated on ice for 1 h. Excess salt and detergent were then removed by passing through PD-10 desalting columns pre-equilibrated with encapsulation buffer. The eluted liposome suspension was then ultra-centrifuged at 100 000 × g, 4 °C for 40 min and the supernatant discarded. The (proteo)liposome pellets were then resuspended to a lipid concentration of 250 mg/mL in assay buffer (20 mM HEPES pH 7.5, 150 mM NaCl, 0.5 mM PIPES and 0.5 mM Tris base). The uptake assay was initiated by addition of 100 µM putrescine, followed by incubation at room temperature. At select time points, 20 µL aliquots of the assay suspension were filtered through Sephadex G-50 (fine) spin columns (Cytiva) pre-equilibrated with base buffer (20 mM HEPES pH 7.5, 150 mM NaCl) by centrifugation. Eluates were snap frozen in liquid nitrogen until LC-MS analysis. For metabolite extraction, liposomes were dissolved in 400 µL 1:1 methanol/chloroform mixture, then 120 µL of water was added to achieve a final methanol/chloroform/water ratio of 5:5:3.

The mixture was vortexed vigorously for 10 s, then centrifuged at 18 000 × g, 4 °C for 10 min. The aqueous supernatant was transferred to HPLC vials, dried under a stream of nitrogen gas, and then stored at -80 °C until LC-MS analysis as described above.

### Flow cytometry

For all flow cytometry experiments, cells were seeded two days prior at an appropriate density such that cells are 80% confluent on the day of collection. Cells were induced +/- 1 µg/mL Dox 24 hours before harvest. On the day of collection, cells are detached using 1 mM EDTA in PBS, washed with PBS, then fixed with 4% formaldehyde in PBS for 10 min. Fixed cells were washed once with PBS, resuspended in PBS + 1% BSA and stored at 4 C for maximum 1 week. For intracellular GNL staining, approximately 3 × 10^5^ cells/well were added to triplicate wells on a V-bottom 96-well plate, centrifuged and supernatant decanted. Cells were permeabilized by resuspension in 100 µL 90% MeOH, 10% PBS and incubation on ice for 30 min. Cells were then aliquoted to V-bottom 96-well plate in triplicate wells, centrifuged at 350 × g for 3 min, supernatant decanted, and blocked by resuspension in PBS + 10% FBS and incubation on a rotary shaker at room temperature for 1 h, followed by primary staining with biotinylated GNL 1:1000 (Vector Labs) in PBS + 10% FBS for 1 h. Cells were washed twice by resuspending in PBS + 10% FBS and incubating with shaking for 5 min each time. Secondary staining was performed with Alexa Fluor 647-conjugated streptavidin for 30 min at room temperature, following which cells were washed twice as above and resuspended in PBS for flow cytometry analysis. For VVL staining, cells were added to V-bottom 96-well plate, centrifuged and supernatant decanted. Cells were then incubated on ice for 30 min with biotinylated VVL (Vector Labs) diluted 1:1000 in TSM buffer, washed by pelleting and resuspension in PBS, secondary stained with Alexa Fluor 488- conjugated streptavidin on ice for 30 min, then wash again and resuspended in PBS. Staining for Tn antigen followed the same procedure, but with 1 h incubation on ice using anti-Tn antibody (5F4 clone) at 1:3 in PBS + 1% BSA and secondary staining with Alexa Fluor 594-conjugated anti-mouse IgM. Flow cytometry analysis was performed on LSR Fortessa (BD Biosciences) flow cytometer with high-throughput microplate sampler.

Flow cytometry data files were exported in FCS 2.0 format and processed in FlowJo (BD Biosciences). For each experiment batch, a first gate was manually set on the forward scatter (FSC)-Area vs side scatter (SSC)-Area surface to isolate non-debris cells. A second gate was then set on the FSC-Area vs FSC-Height surface to isolate single cells. Fluorescence values for individual gated cells were exported and analyzed in R. Since value × g varied between cell populations and were often non-Gaussian and/or bimodal, Cliff’s delta was calculated using the *effsize* package and used as a non-parametric measure of effect size between Dox/uninduced samples of each cell line.

### Immunofluorescence

HCT116 SLC45A4 KO-OE or HEK293 MFSD5 WT-OE cells were seeded at 150 000 cells/well on polylysine-coated glass cover slips placed in 24-well plates and induced with 1 µg/mL Dox overnight. Media was aspirated and cells were fixed in 4% formaldehyde in PBS for 10 min at room temperature. Cells were then blocked and permeabilized with IF blocking buffer (PBS + 10% FBS + 0.3% Triton X-100) for 1 h, then primary antibodies or lectins added at the following dilutions in IF blocking buffer: rat anti-HA 1:1000 (Roche), biotinylated GNL 1:1000 (Vector Labs), mouse M6P-1 1:50 (ABCD Antibodies), rabbit anti-LAMP1 1:200 (Cell Signaling). Following two hours of gentle rocking at room temperature, cells were washed three times with IF blocking buffer, then secondary staining performed for 1 h with the following antibodies: anti-mouse Alexa Fluor 568 1:1000, streptavidin Alexa Fluor 647 1:1000, anti-rabbit Alexa Fluor 488 1:1000, DAPI 1:2000. Cells were washed three times, mounted on glass slides with Fluoromount-G (Thermo Fisher), then imaged with an LSM 700 confocal microscope (Zeiss).

### Glycoproteomics

Cells were seeded two days prior to harvest at an appropriate density such that cells are 80% confluent on the day of collection. Cells were induced +/- 1 µg/mL Dox 24 hours before collection. On the day of harvest, cells are detached using 1 mM EDTA in PBS, washed twice with PBS to remove residual EDTA, then pelleted and stored at -80 C until lysis. Cell pellets were lysed using a buffer containing 50 mM Tris-HCl (Trizma Base: Sigma-Aldrich, Cat#T6066; HCl: Sigma-Aldrich, Cat#258148), 8M urea (VWR, Cat#0568), 150 mM NaCl (Merck, Cat#106404), pH 8, containing phosphatase inhibitors (Cell Signaling Technology, Cat#5870). Samples were kept on ice for 30 minutes before being sonicated using a water bath sonicator for 1 minute at low intensity. Samples were centrifuged at 20 000g for 15 min at 4 °C. Resulting supernatants were transferred to a new tube, and protein contents were quantified using Pierce™ BCA Protein Assay Kit (Thermo Fischer, Cat#23225). For each sample, 300 µg of protein were reduced using DTT (Roche, Cat#10708984001; final concentration 10 mM) for 30 min at 37 °C, and subsequently alkylated using IAA (Sigma-Aldrich, Cat#I6125; final concentration 20 mM) for 30 min in the dark at room temperature. Samples were then diluted using a 50 mM Tris-HCl buffer to 4M urea before adding 6 µg of Lys C (FUJIFILM Wako, Cat#125 05061) to each sample (1 µg Lys C: 50 µg sample protein). Samples were incubated for 2 h at 30 °C, before being further diluted by adding 50 mM Tris-HCl buffer to lower the urea concentration to 2M. 6 µg of Trypsin Gold (Promega, Cat#V5280) were added to each sample (1 µg Trypsin: 50 µg sample protein), and samples were further incubated for 15 h at 37 °C. At this point, samples were acidified using TFA to a final concentration of 0.5% before being cleaned using 200 mg (3cc) Sep-Pak cartridges (Waters, Cat#WAT054945). The resulting elution fraction containing the cleaned peptides was dried under vacuum and stored at -20 °C. For TMT labelling, 200 µg of peptides of each sample were prepared in a final volume of 100 µL of 100 mM HEPES buffer pH 7.6. Samples were labeled using TMTpro 18 plex (Thermo Scientific, Cat#A52045) reagents, according to manufacturer’s instructions. The labelling efficiency was determined by LC-MS/MS on a small aliquot of each sample. After quenching the labeling reaction, samples were mixed in equimolar amounts, evaluated by LC-MS/MS. The mixed sample was acidified to a pH below 2 with 10% TFA and desalted using C18 cartridges (Sep-Pak Vac 1cc (200mg), Waters, Cat#WAT054945). Peptides were eluted with 2 x 600 μl 80% Acetonitrile (ACN) and 0.1% Formic Acid (FA), followed by freeze-drying. Glycopeptides from the resulting TMTpro mixes were enriched by performing off-line ion pairing (IP) HILIC chromatography, as described previously (12). Briefly, the dried samples were taken up in 100 μL 75% acetonitrile containing 0.1% TFA and subjected to chromatographic separation on a TSKgel Amide-80 column (4.6 x 250 mm, particle size 5μ) using a linear gradient from 0.1% TFA in 80% acetonitrile to 0.1% TFA in 40% acetonitrile over 35 min (Dionex Ultimate 3000, Thermo). The 30 collected fractions were vacuum dried. Samples were resuspended using 0.1% trifluoroacetic acid (TFA, Thermo Scientific, Cat#28903). The IP-HILIC fractions were individually analysed by LC–MS/MS. The nano HPLC system used was an UltiMate 3000 HPLC RSLC nano system (Thermo Scientific) coupled to a Q Exactive HF-X mass spectrometer (Thermo Scientific), equipped with a Proxeon nanospray source (Thermo Scientific). Peptides were loaded onto a trap column (Thermo Scientific, PepMap C18, 5 mm × 300 μm ID, 5 μm particles, 100 Å pore size) at a flow rate of 25 μL/min using 0.1% TFA as mobile phase. After 10 min, the trap column was switched in line with an analytical column (Thermo Scientific, PepMap C18, 500 mm × 75 μm ID, 2 μm, 100 Å). Peptides were eluted using a flow rate of 230 nl/min and a binary 180 min gradient. The two steps gradient started with the mobile phases: 98% A solution (water/formic acid, 99.9/0.1, v/v) and 2% B solution (water/acetonitrile/formic acid, 19.92/80/0.08, v/v/v), which was then increased to 35% B over the next 180 min, followed by a gradient of 90% B for 5 min, which was finally, in a 2 min period, decreased to the gradient 95% A and 2% B for equilibration at 30 °C. The following parameters were used for MS acquisition using an Exploris 480 instrument, operated in data-dependent mode: the instrument was operated in a positive mode; compensation voltages (CVs) used = CV-40, CV-50, CV-60; cycle time (per CV) = 1 sec. The monoisotopic precursor selection (MIPS) mode was set to Peptide. Precursor isotopes and single charge state precursors were excluded. MS1 resolution = 60 000; MS1 Normalised AGC target = 300%; MS1 maximum inject time = 50 msec. MS1 restrictions were relaxed when few precursors were present. MS1 scan range = 400–1,600 m/z; MS2 resolution = 45 000; MS2 Normalised AGC target = 200%. Maximum inject time = 250 msec. Precursor ions charge states allowed = 2-7; isolation window = 1.4 m/z; fixed first mass = 110 m/z; HCD collision energies = 30,33,36; exclude isotopes = True; dynamic exclusion = 45 s. All MS/MS data were processed and analyzed using Xcalibur v3.1 (Thermo), and Byonic (Protein Metrics) included in Proteome Discoverer v3.0. Two default glycan databases (Mammalian O-glycans and Human N-glycans) were used with the latest human or mouse UniProt database, to generate the glycopeptide search space. We used an in-house developed R script to filter out poorly matching spectra, considering a Byonic score higher than 200. Significant (1% and 5% FDR) changes in glycopeptide abundances were determined by considering p-value and magnitude of fold change simultaneously as previously described (Hein *et al*, 2015). Further data processing, analysis and visualization were performed in R.

## Data availability

The data is available through the RESOLUTE web portal and corresponding dashboards (https://re-solute.eu/resources). Raw data are submitted to MetaboLights and European Nucleotide Archive (ENA).

## Acknowledgements

This study received funding from the RESOLUTE consortium. RESOLUTE has received funding from the Innovative Medicines Initiative 2 Joint Undertaking under grant agreement No 777372. This Joint Undertaking receives support from the European Union’s Horizon 2020 research and innovation programme and EFPIA.

D.B.S. and G.C. are supported by the Ontario Institute for Cancer Research, Royal Institution for the Advancement of Learning McGill University, Kungliga Tekniska Hoegskolan, Diamond Light Source Limited and by the Innovative Medicines Initiative 2 Joint Undertaking (JU) under grant agreement No 875510. The JU receives support from the European Union’s Horizon 2020 research and innovation program and EFPIA.

The lab of J.M.P. received funding from the Medical University of Vienna, the T. von Zastrow foundation. S.M. was supported by the European Union’s Horizon 2020 research and innovation programme under the Marie Sklodowska-Curie grant agreement No 841319 and the ESPRIT-Programme of the Austrian Science Fund (FWF, Project number: ESP 166).

G.S-F. was supported by the Austrian Academy of Sciences.

We thank U. Mandel and H. Clausen (University of Copenhagen) for providing 5F4 hybridoma supernatant.

We highly appreciate the feedback and critical reading of the manuscript by Ann-Katrin Hopp, Enrico Girardi, Gabriel Onea, Kai-Chun Li, Leonhard Heinz, Manuele Rebsamen, and Vojtech Dvorak.

This article reflects only the authors’ views and neither IMI nor the European Union and EFPIA are responsible for any use that may be made of the information contained therein.

## Conflict of interest

G.S-F. is co-founder and owns shares of Solgate GmbH, an SLC-focused company.

## Tables and legends

Table 1: Cell lines

Table 2: Targeted metabolomics data set (normalized and batch-corrected peak areas)

Table 3: Differential metabolite abundance and categorization of SLC-metabolites pairs

Table 4: Pathway coverage and regulation

Table 5: Differential expression analyses underlying uniqueness analysis

Table 6: Standard normalized differential metabolite abundance for clustering

Table 7: Standard normalized differential expression for clustering

Table 8: Reactome pathway average gene expression of transcriptome clusters

Table 9: Glycosylation-related cluster SLCs effects on intracellular GNL staining

Table 10: Glycosylation-related cluster SLCs effects on cell surface VVL staining

Table 11: Targeted metabolomics analysis of culture media

Table 12: Targeted metabolomics method (internal standard assignment, MRM transition, retention times)

## Figures and Expanded View Figures

**Figure EV1.**
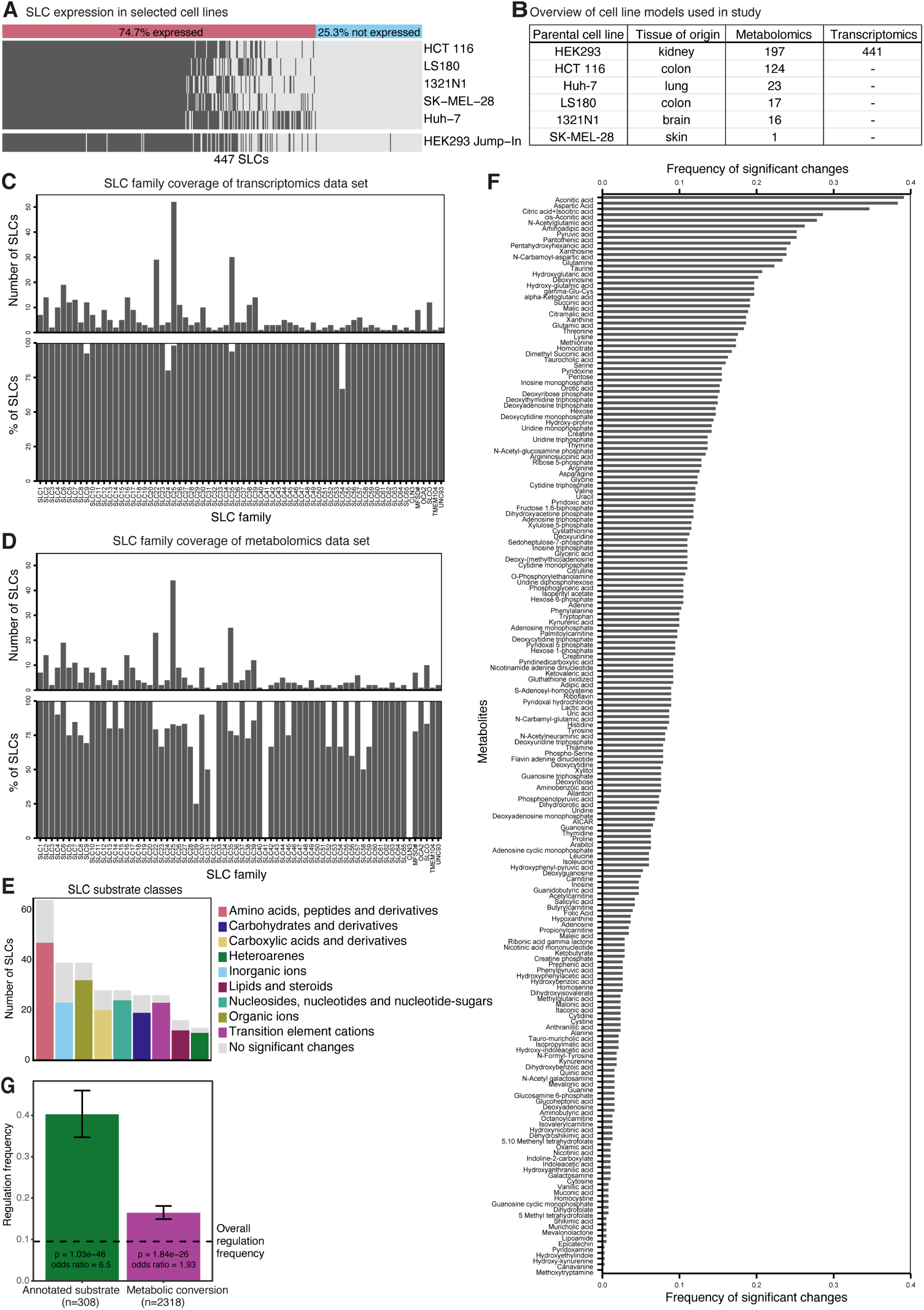
Comprehensive multi-omic coverage of the SLC superfamily. **(A)** Heatmap illustrating coverage of 447 SLCs across five cancer cell lines considered for KO and KO-OE (HCT 116, LS180, 1321N1, SK-MEL-28, Huh-7) as well as Jump-In™ T-REx™ HEK293. Tpm > 1 was used as threshold for expression. **(B)** Number of SLCs with targeted metabolomics and transcriptomics data sets and respective parental cell lines included in this study. **(C)** Coverage of transcriptomics data sets in this study compared to the SLC superfamily (the RESOLUTE list of 447 SLCs). The coverage is given both in absolute numbers per family (top) and percentage of members per family (bottom). **(D)** Coverage of targeted metabolomics data sets in this study compared to the SLC superfamily (the RESOLUTE list of 447 SLCs). The coverage is given both in absolute numbers per family (top) and percentage of members per family (bottom). **(E)** Profiled SLCs by targeted metabolomics and divided according to the substrate class of their annotated substrates. Colored bars indicate proportion of SLCs with significant changes upon differential analysis of metabolite abundance +/- doxycycline (dox) induction, while grey bars indicate proportion of SLCs without significant metabolite changes. **(F)** Frequency of significant changes per metabolite across all profiled cell lines. Frequency was calculated for each of the 189 metabolites as the fraction of cell lines with its significant change from the total of 378 profiled SLCs. **(G)** Frequency of significant changes of annotated substrates/metabolic conversions vs overall regulation frequency. All pairs of SLCs and targeted metabolites were grouped by a potential match of the SLC’s annotated substrates to the targeted metabolite. The frequency of differential regulation was calculated per group and was found to be significantly higher for the 308 cases where a targeted metabolite could be directly matched to an annotated substrate for the overexpressed SLC. A smaller but also significant effect was found for the 2318 cases where a targeted metabolite could be matched via metabolic conversion to an annotated substrate for the overexpressed SLC (both Fisher’s exact tests; error bars are calculated from the odds ratio confidence region).

**Figure EV2.**
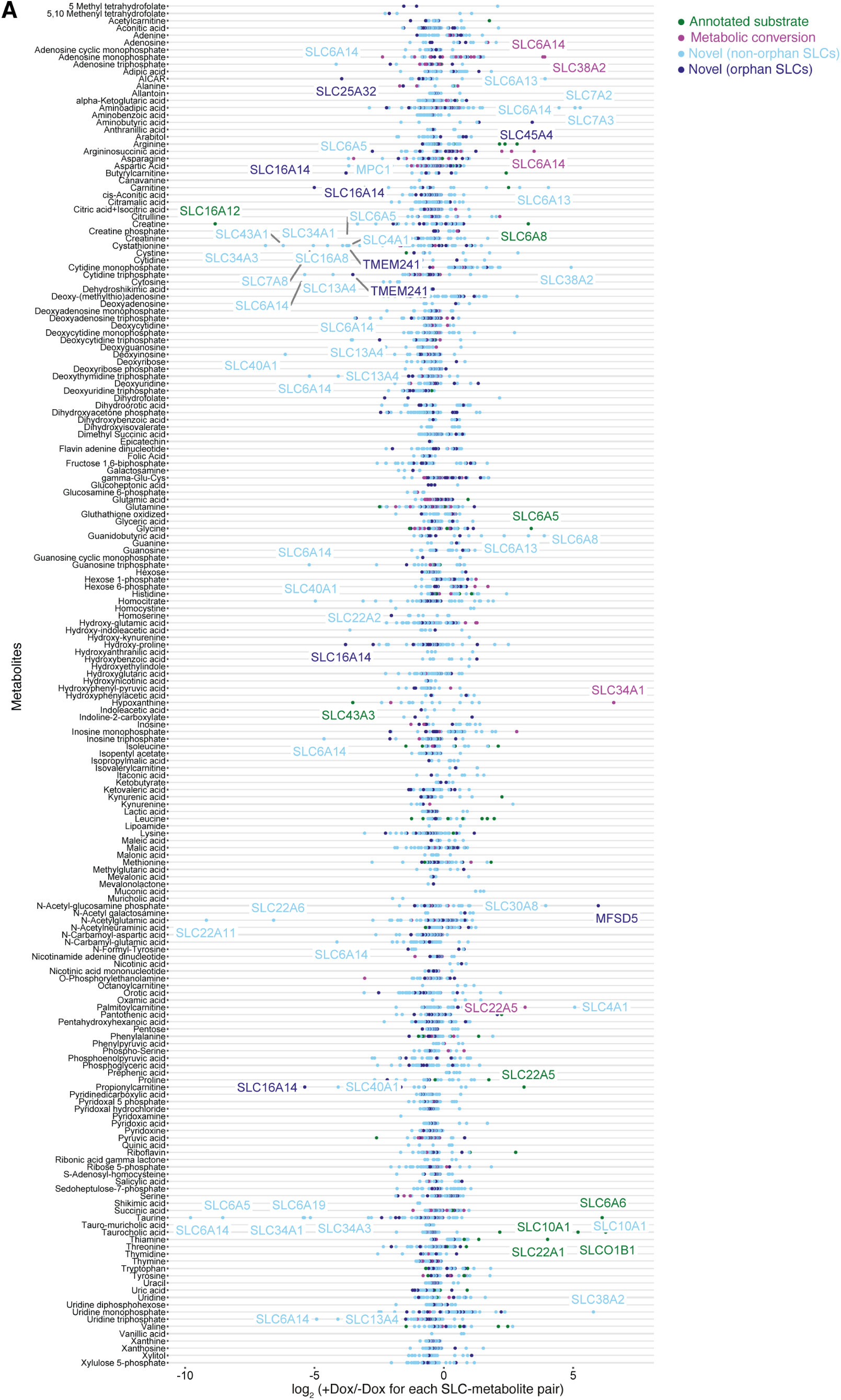
Overview of SLC-metabolite pairs by category and per metabolite. **(A)** Visualization of all significant SLC- metabolite pairs (p-adj. < 0.05) in a per metabolite view with metabolites on the y-axis and log2 +/-doxycycline (dox) for each SLC-metabolite pair on the x-axis. Colors indicate the SLC-metabolite pair category (‘annotated substrate’, ‘metabolic conversion’, ‘novel (non-orphan SLCs)’, ‘novel (orphan SLCs)’.

**Figure EV3.**
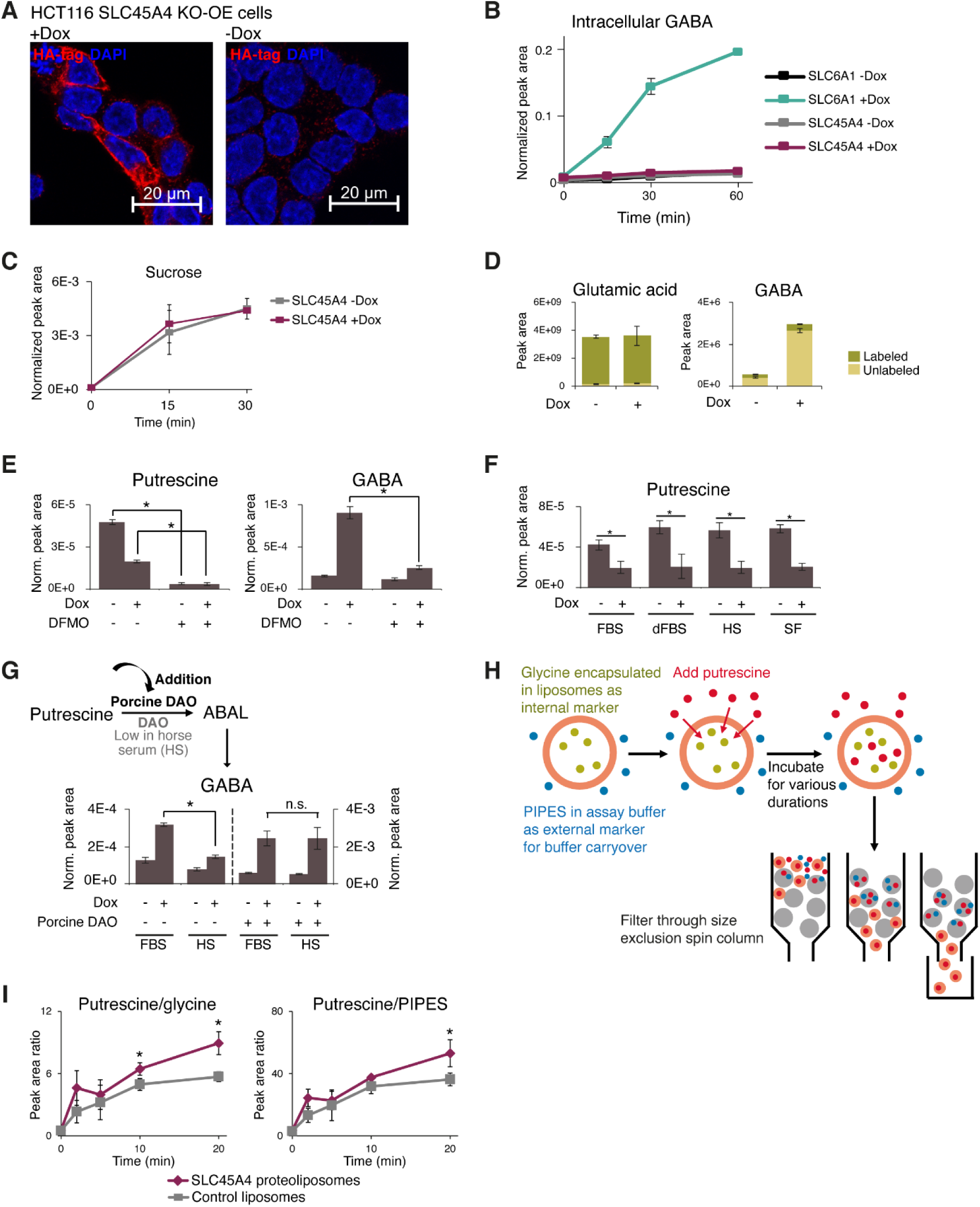
SLC45A4 mediates GABA production via putrescine export and oxidation. **(A)** Representative immunofluorescence images of HCT116 SLC45A4 KO-OE cells +/- 24 h doxycycline (Dox) induction. Red and blue channels show HA-tagged SLC45A4 and DAPI nuclear counterstain, respectively. **(B)** Intracellular GABA levels in HCT116 Renilla KO cells transduced with Dox-inducible SLC6A1 overexpression construct and HCT116 SLC45A4 KO-OE cells, following +/- 24 h Dox induction in OptiMEM media +dFBS and switch to OptiMEM +dFBS +100 µM GABA for the indicated durations. Data points represent means, error bars represent s.d. (n=3). **(C)** Intracellular sucrose levels in HCT116 SLC45A4 KO-OE cells following +/- 24 h Dox induction and switch to media supplemented with 1 mM sucrose for the indicated durations. Data points represent means, error bars represent s.d. (n=3). **(D)** Labeled (from ^13^C^15^N-glutamic acid) and unlabeled abundances of intracellular metabolites in HCT116 SLC45A4 KO-OE cells +/- 24 h Dox induction. Bar heights represent means, error bars represent s.d. (n=6). **(E)** Intracellular putrescine and GABA levels in HCT116 SLC45A4 KO-OE cells with and without ODC1 inhibition by 1 mM difluoromethylornithine (DFMO) +/- 9 h Dox induction. Bar heights represent means, error bars represent s.d. (n=3). **(F)** Intracellular putrescine levels in HCT116 SLC45A4 KO-OE cells cultured in regular FBS-containing media, media with dialyzed FBS (dFBS), media with horse serum (HS), or serum-free media (SF) +/- 9 h Dox induction. Bar heights represent means, error bars represent s.d. (n=3). **(G)** Intracellular GABA levels in HCT116 SLC45A4 KO-OE cells cultured in regular FBS-containing media vs horse serum (HS)-containing media, without (left) or with (right) porcine diamine oxidase (DAO) supplementation +/- 9 h Dox induction. Bar heights represent means, error bars represent s.d. (n=3). **(H)** Schematic of cell-free putrescine uptake assay. Glycine is encapsulated within liposomes inserted with SLC45A4 protein or protein-free control liposomes as an internal marker of liposome abundance. Liposomes are resuspended in assay buffer containing PIPES as an external marker of buffer carryover. Following addition of 100 µM putrescine and incubation for defined time points, liposome suspensions are filtered through Sephadex G50 size exclusion chromatography (SEC) spin columns, which traps assay buffer components while allowing liposomes to flow through. The liposome-enriched eluates are then analyzed by LC-MS. **(I)** Putrescine/glycine and putrescine/PIPES peak ratios in SEC eluates of SLC45A4 proteoliposomes vs control liposomes over different uptake durations. Data points represent means, error bars represent s.d. (n=3). Where presented, asterisks (*) and n.s. on significance bars indicate p < 0.05 and p ≥ 0.05 (Student’s t-test) respectively.

**Figure EV4.**
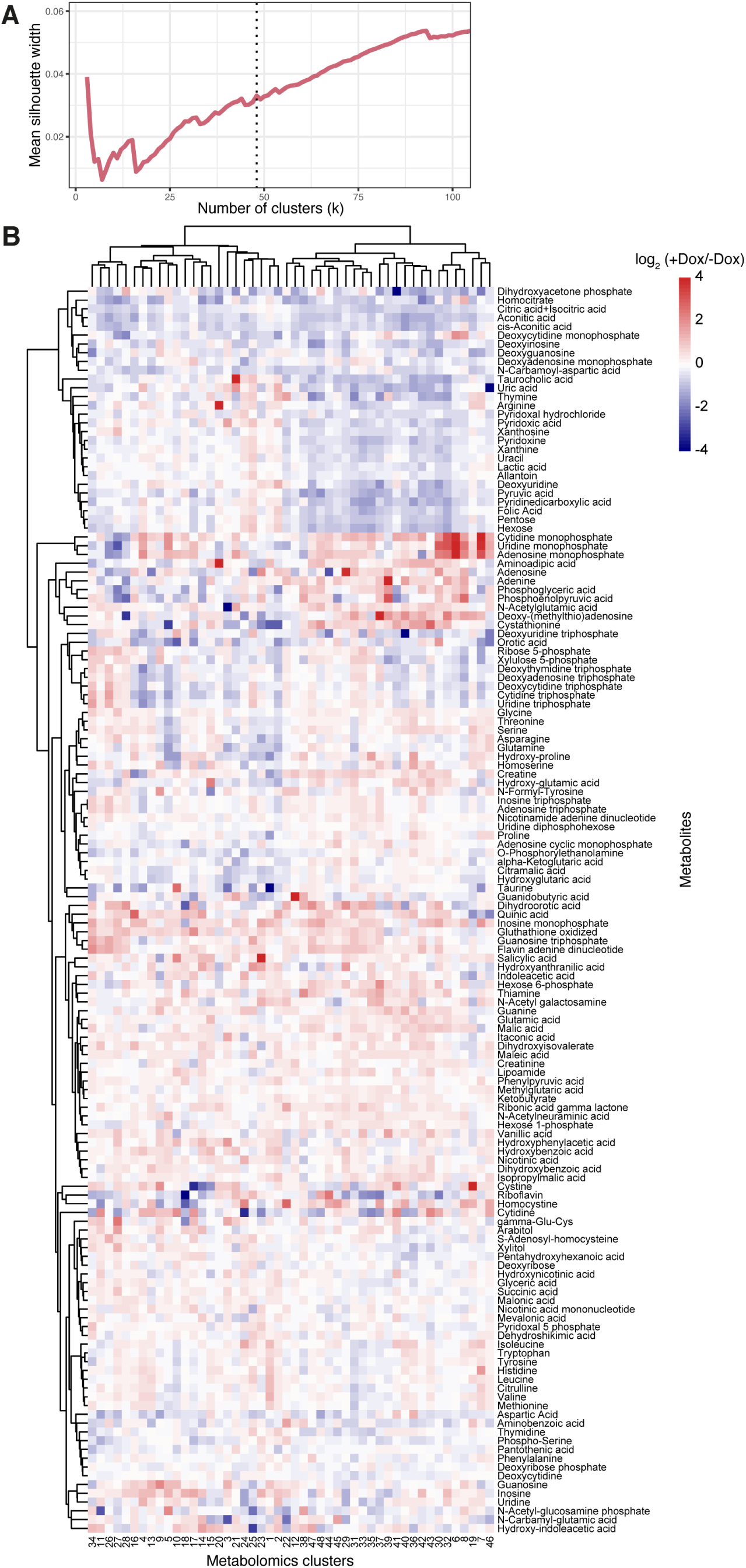
Metabolite-centric analysis of similarity clusters reveals predominantly deregulated and co-regulated metabolites. **(A)** Mean silhouette width across all clusters at different cluster numbers (k=2-100). The dashed line indicates the chosen k=48. **(B)** Heatmap of all clusters and metabolites based on the respective averaged log2 fold changes per cluster. Cluster numbers are indicated on x-axis. Metabolites are indicated on y-axis.

**Figure EV5.**
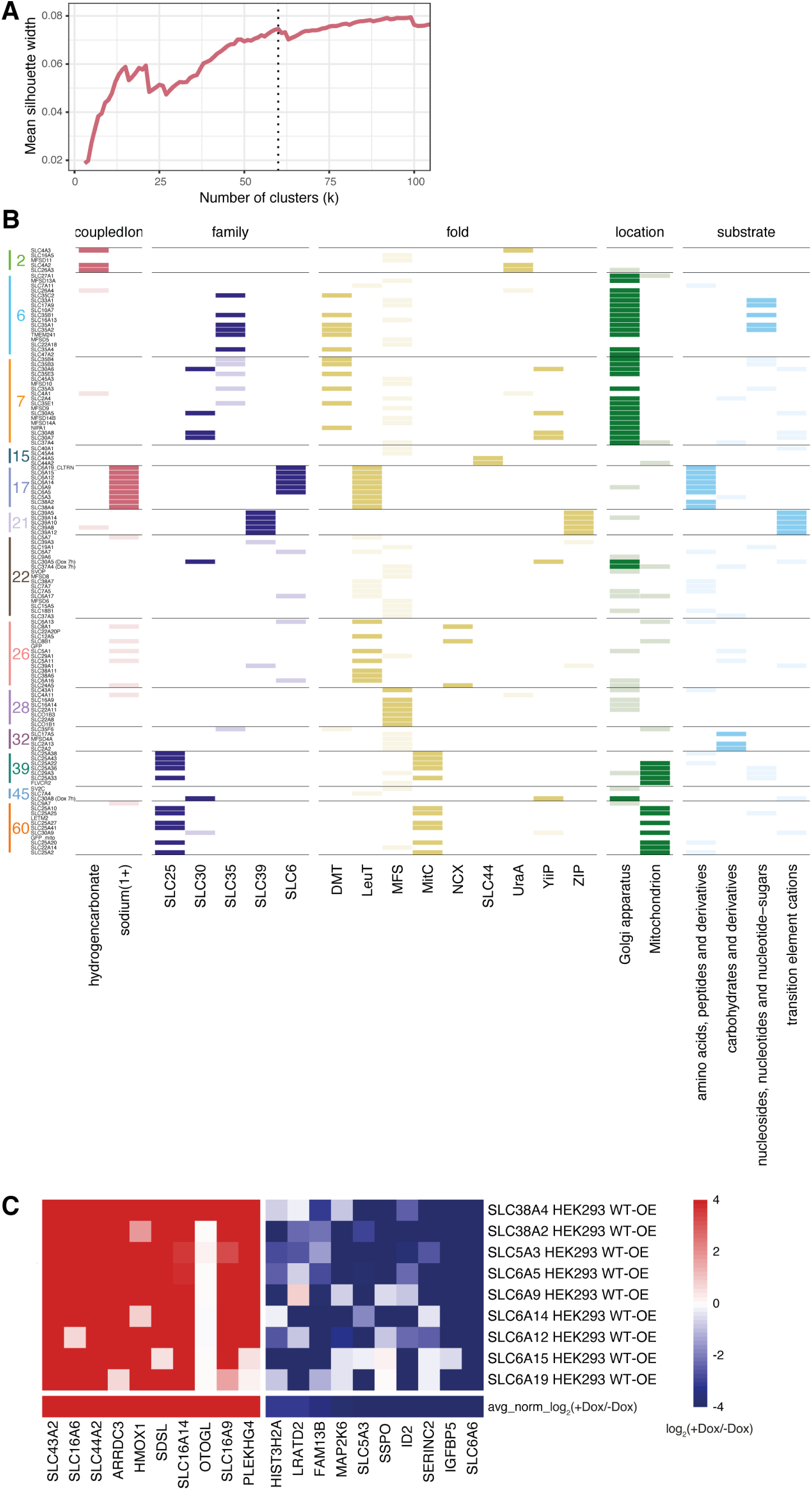
Enrichment of annotated functional SLC features in transcriptomic clusters. **(A)** Mean silhouette width across all clusters at different cluster numbers (k = 2–100). The dashed line indicates the chosen k=60. **(B)** Transcriptomic clusters with significant SLC functional feature enrichment (Fisher’s test p < 0.2). **(C)** Heatmap showing doxycycline (Dox) vs uninduced log2 fold changes of the 20 most deregulated genes in cluster 17 for each cluster member. Genes are indicated on x-axis and the clustered cell lines on the y-axis.

**Figure EV6.**
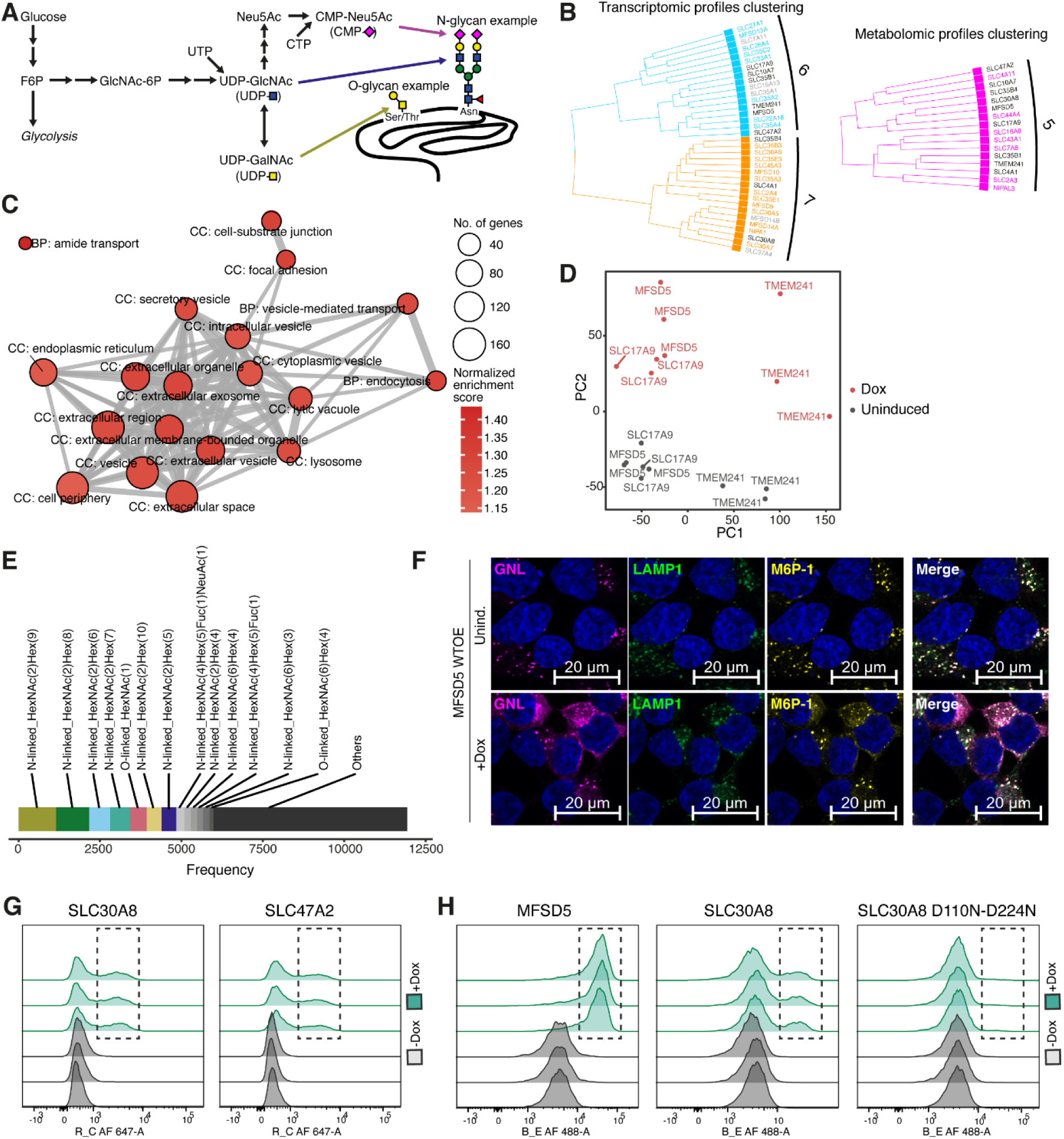
SLC cluster alters both N-and O-linked glycosylation signatures. **(A)** Schematic representation of hexosamine and sialic acid biosynthetic pathway that produces nucleotide sugar substrates for glycosyltransferase reactions. **(B)** Transcriptomics clusters 6 and 7 (left) and metabolomics cluster 5 (right) showing common SLCs between clusters from the two different data modalities. **(C)** GO terms significantly enriched (p-adj. < 0.05) by gene set enrichment analysis (GSEA) on most abundant glycoproteins detected in the data set. Abundances were calculated as total signal intensity of corresponding glycopeptides for each protein. **(D)** Principal components analysis performed on normalized glycopeptide abundances of 18 glycoproteomics samples, whereby to account for possible differences in protein amount, each glycopeptide was normalized to the total abundance of the corresponding protein. **(E)** Stacked bar plot indicating number of glycopeptides attributed to each glycan composition, showing that 7 most abundant glycan compositions account for ∼40.6% (4845 out of 11938) identified glycopeptides. **(F)** Representative immunofluorescence images of MFSD5 WT-OE uninduced and Doxycycline (Dox)- induced cells showing individual and merged channels for *Galanthus Nivalis* Lectin (GNL), LAMP1 and M6P staining with DAPI nuclear counterstain. **(G)** Flow cytometry histograms comparing cell surface VVL staining of SLC30A8 and SLC47A2 WT-OE +/- Dox cells. Dashed boxes indicate +Dox cell populations with increased VVL staining. **(H)** Flow cytometry histograms comparing cell surface VVL staining of MFSD5, SLC30A8 wild-type, and SLC30A8 D110N-D224N (transport deficient mutant) +/- Dox cells. Dashed boxes indicate +Dox cell populations with increased VVL staining, which is absent in the transport deficient mutant.

